# Responses to conflicting binocular stimuli in mouse primary visual cortex

**DOI:** 10.1101/2024.12.31.630912

**Authors:** Daniel P. Montgomery, Daniel A. Bowen, Jin Wu, Mark F. Bear, Eric D. Gaier

**Affiliations:** Center for Neuroscience Research, Children’s National Hospital, Washington, DC; Department of Ophthalmology, Boston Children’s Hospital, Boston, MA; Picower Institute for Learning and Memory, Massachusetts Institute of Technology, Cambridge, MA; Brandeis Neuroscience Graduate Program, Brandeis University, Waltham, MA; Harvard Medical School, Boston, MA

**Keywords:** Binocular vision, interocular suppression, binocular rivalry

## Abstract

Binocular vision requires that the brain integrate input from both eyes to form a unified percept. Small interocular differences support depth perception (stereopsis), while larger disparities can cause diplopia or binocular rivalry. The neural mechanisms by which early visual circuits process concordant versus conflicting binocular signals remain incompletely understood, particularly in the case of large disparities. Here, we used visually evoked potential (VEP) recordings, unit recordings, and 2-photon calcium imaging in the binocular region of mouse primary visual cortex (bV1) to examine how distinct forms of binocular disparity engage local circuits. Using a dichoptic display, we found that interocular phase disparities reduced VEP magnitude through decreased neuronal firing early in the response (40-80 ms after stimulus onset). In contrast, orientation disparities also decreased VEP magnitude, but via increased firing later in the response (100-200 ms). This late activity was enhanced in both regular-spiking (putative excitatory) and fast-spiking (putative parvalbumin-positive inhibitory) units. In contrast, calcium imaging revealed that somatostatin-positive interneurons were suppressed during orientation conflict. These findings suggest that phase differences suppress bV1 responses via feedforward mechanisms, while orientation disparities prolong activity through disinhibition mediated by somatostatin-positive interneurons. Our results identify distinct circuit pathways recruited by different forms of binocular conflict, clarify how early visual cortex contributes to binocular integration, and provide a foundation for investigating perceptual suppression and rivalry.

**IMPACT STATEMENT:** Distinct forms of binocular conflict engage separate circuit mechanisms in mouse primary visual cortex, revealing how interocular disparities shape population activity through feedforward and disinhibitory processes.

## INTRODUCTION

Visual processing of information coming from the two eyes requires the integration of conflicting binocular signals. Under physiologic conditions, disparities between the images projected onto each retina tend to be modest, and the brain can effectively fuse the two images into a single percept. Processing of spatial image disparities, within a range, enables the perception of depth (stereopsis) (Parker 2007). How the brain processes conflicting visual signals has been studied since the foundational work of Hubel and Wiesel (Hubel and Wiesel 1962). In mammals, signals from the two eyes largely first converge in primary visual cortex (V1) where monocularly responsive neurons with retinotopically aligned receptive fields synapse onto binocular neurons. Many binocular neurons are modulated by modest spatial differences between the inputs from each eye, a phenomenon known as disparity-tuning. Disparity-tuned neurons are thought to serve stereopsis (Hubel and Wiesel 1970, Hubel and Livingstone 1987, Cumming and DeAngelis 2001). In mice, even relatively large spatial disparities can be integrated by neurons in V1 (Scholl, Burge et al. 2013, Scholl, Pattadkal et al. 2017, La Chioma, Bonhoeffer et al. 2020), corresponding to the short viewing distance at which mice tend to interact with objects (Samonds, Choi et al. 2019).

Although disparity-tuned neurons in V1 have been extensively studied under conditions of modest spatial offset, less is known about how cortical circuits respond when binocular input is strongly conflicting. In strabismus, a clinical condition defined by eye misalignment, large interocular disparities exceed the capacity for binocular integration and preclude stereopsis (Harrad, Sengpiel et al. 1996). When disparities between the images viewed by each eye are too great, fusion breaks down into diplopia (double vision) and binocular rivalry (an alternating, bistable percept). These outcomes can arise from a range of interocular differences in visual input, suggesting that different types of conflict may engage distinct cortical mechanisms. However, how early visual circuits respond to qualitatively different forms of interocular conflict remains poorly understood.

We sought to study how mammalian V1 processes large interocular image disparities using the tractability of the mouse visual system to identify the populations of neurons involved. Our goal was to identify how distinct forms of interocular conflict are represented at the level of local circuits in primary visual cortex, providing a critical step toward understanding the neural basis of perceptual suppression. Using a combination of visually evoked potentials (VEPs), unit recordings, and 2-photon (2P) calcium imaging, we characterized cortical responses to two types of interocular stimulus disparities in mice. The first—orientation-matched grating stimuli with an interocular phase difference—mimics the magnitude of spatial disparity that tends to give rise to diplopia in humans. The second—orthogonally oriented grating stimuli— employs stimulus combinations often used to study binocular rivalry in humans (Blake and Logothetis 2002) and non-human primates (Leopold and Logothetis 1996). Although both types of disparity led to reductions in VEP magnitude, they did so via markedly different mechanisms: phase offset stimuli suppressed the early negative component of the VEP through decreased early evoked firing, while orthogonal stimuli suppressed the late positive component through prolonged spiking activity. These stimulus-specific patterns were evident across cortical layers and in the responses of both regular-spiking (RS, putative excitatory) and fast-spiking (FS, putative parvalbumin-positive inhibitory) neurons. Furthermore, 2P calcium imaging revealed that a separate class of interneurons expressing somatostatin (SOM+) exhibited decreased activity in response to orthogonal stimuli. These findings demonstrate that V1 circuits in mice exhibit distinct patterns of activity in response to different forms of interocular conflict. This divergence is reflected in anatomically and temporally specific activity patterns across excitatory and inhibitory cell types, providing mechanistic insight into how early visual cortex processes discordant binocular input.

## RESULTS

### Interocular phase or orientation disparities differentially reduce VEP magnitude

We first sought to determine the effect of presenting disparate stimuli to each eye on the cortical responses of mice. We implanted mice with tungsten microelectrodes in layer 4 (L4) of bV1 (**Figure 1A**) and measured VEPs in awake, head-fixed mice using phase-reversing sinusoidal grating stimuli presented on a passive 3D display (**Figure 1B**). Reciprocally polarized lenses placed in front of each eye enabled the independent presentation of binocularly discordant stimuli. For all recordings, the eye contralateral to the recording electrode viewed full contrast stimuli at a 45° orientation. The ipsilateral eye viewed stimuli ranging from 0% contrast (i.e. grey screen) to 50% contrast. Ipsilateral eye stimuli were either in phase (concordant) or 180° out of phase (phase offset) relative to the contralateral eye stimulus (**Figure 1C**). We found that the VEP magnitudes (peak negative to peak positive voltage) elicited by concordant stimuli were larger than those elicited by monocular (0% ipsilateral eye contrast) or phase offset stimuli (**Figure 1D**). Notably, a statistically significant difference between concordant and phase offset stimuli emerged at an ipsilateral eye contrast as low as 20%.

**Figure 1.**
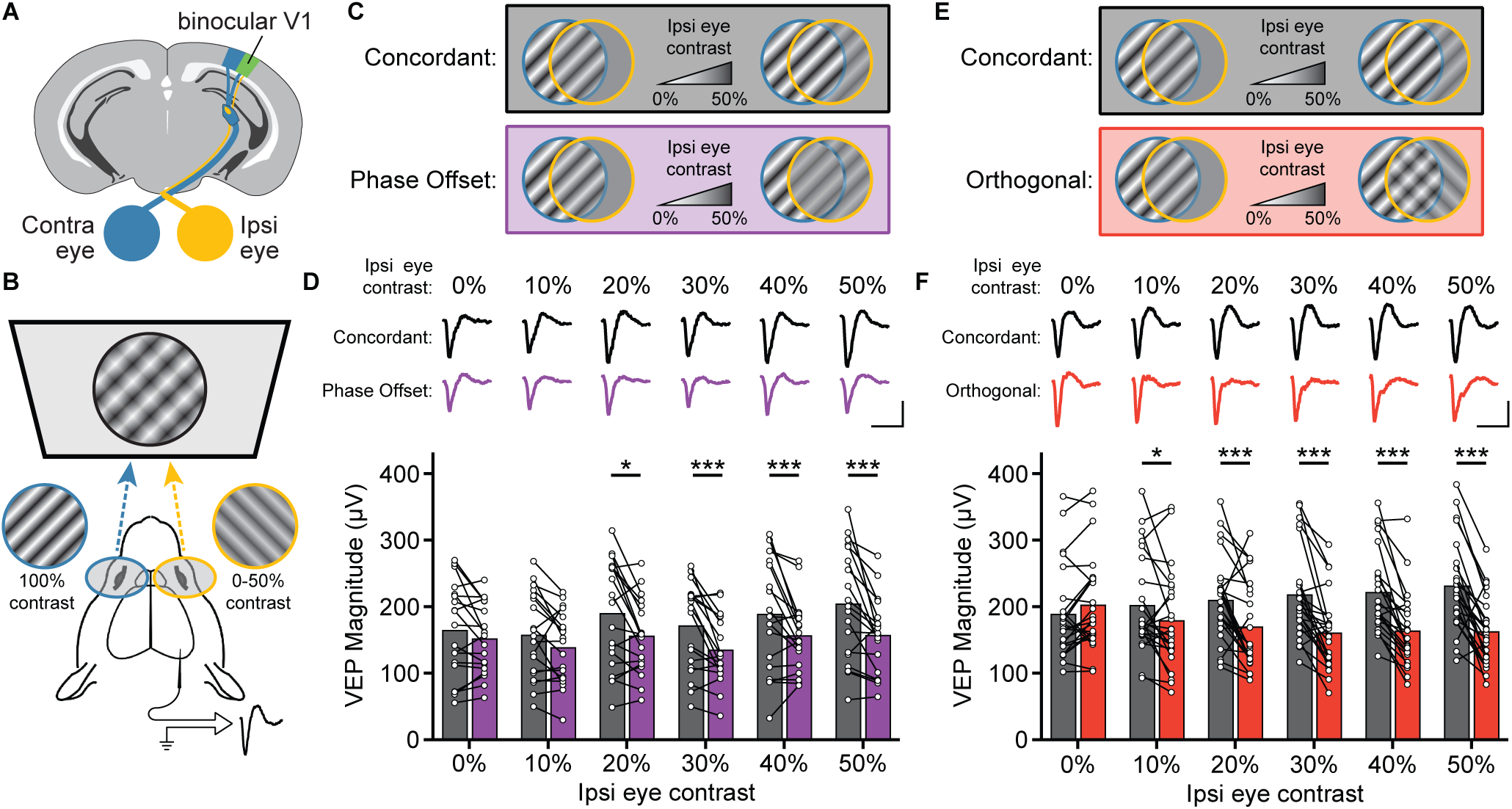
Binocular stimuli with an interocular phase or orientation disparity elicit smaller visually evoked potentials. **(A)** Schematic of mouse brain showing placement of a chronic recording electrode in binocular V1 (V1b). **(B)** Mice viewed phase reversing sinusoidal grating stimuli on a dichoptic display. **(C)** Phase offset experimental design. The contralateral (Contra) eye viewed full-contrast stimuli, while the ipsilateral (Ipsi) eye viewed stimuli ranging from 0% to 50% contrast, presented either in phase (Concordant) or out of phase (Phase Offset) with the contra eye stimulus. **(D)** For ispi eye contrasts ≥20% contrast, concordant stimuli elicited significantly larger VEPs than phase offset stimuli. Average VEP waveforms for each condition shown above bars. Two-way repeated measures ANOVA (main effect of condition, F(1,18) = 16.4, p<0.001; condition × contrast, F(5,90) = 2.37, p = 0.045) followed by Šídák’s multiple comparisons test (concordant vs. phase offset at 0%, 10%, 20%, 30%, 40%, and 50% ipsi eye contrast: p = 0.522, 0.126, <0.001, <0.001, <0.001, <0.001; Concordant 0% vs. 10%, 20%, 30%, 40%, and 50% contrast: p = 0.999, 0.088, 0.999, 0.110, <0.001; Phase Offset 0% vs. 10%, 20%, 30%, 40%, and 50% contrast: p = 0.955, 0.999, 0.678, 0.999, 0.999). **(E)** Orthogonal orientation experimental design. Similar to **C**, but the ipsi eye viewed stimuli at either the same orientation as the contra eye (Concordant) or rotated 90° (Orthogonal). **(F)** For ispi eye contrasts ≥10% contrast, Concordant stimuli elicited larger VEPs than Orthogonal stimuli. Two-way repeated measures ANOVA (main effect of condition, F(1,23) = 32.6, p<0.001; condition × contrast, F(5,115) = 13.8, p < 0.001) followed by Šídák’s multiple comparisons test (Concordant vs. Orthogonal at 0%, 10%, 20%, 30%, 40%, and 50% ipsi eye contrast: p = 0.437, 0.037, <0.001, <0.001, <0.001, <0.001; Concordant 0% vs. 10%, 20%, 30%, 40%, and 50% contrast: p = 0.968, 0.282, 0.017, 0.003, <0.001; Orthogonal 0% vs. 10%, 20%, 30%, 40%, and 50% contrast: p = 0.137, 0.003, <0.001, <0.001, <0.001). Scale bars: 200 ms and 100 µV.

Next, we looked at how orientation differences between the two eyes affects bV1 responses. To test this, we used the same dichoptic display to present stimuli that were either at the same orientation for each eye (concordant) or rotated 90° (orthogonal). As before, the contralateral eye viewed a full contrast stimulus while the ipsilateral eye viewed stimuli ranging from 0% to 50% contrast (**Figure 1E**). Again, a substantial contrast-dependent difference in VEP magnitude between concordant and orthogonal ipsilateral eye stimuli emerged, with a statistically significant difference between concordant and orthogonal stimuli at just 10% ipsilateral eye contrast (**Figure 1F**). Relative to monocular stimuli, we also observed a robust increase in VEP magnitude for concordant stimuli, and decrease in VEP magnitude for orthogonal stimuli. These findings suggest that even low contrast engagement of the inherently weaker ipsilateral eye is sufficient to modulate cortical responses in the context of interocular stimulus disparities, particularly in the case of rivalry.

To better understand which features of the VEP were modulated by these disparities, we separately analyzed the negative and positive components of the waveform. For phase offset stimuli, the reduction in VEP magnitude was primarily due to a significant attenuation of the negative peak (**Figure 2A**), while changes in the positive peak were smaller and more variable (**Figure 2B**). In contrast, orthogonal stimuli led to a substantial reduction in the positive component of the VEP, with more modest effects on the negative component (**Figure 2C-D).** These results suggest that distinct features of the VEP are differentially modulated by interocular phase and orientation disparities.

**Figure 2.**
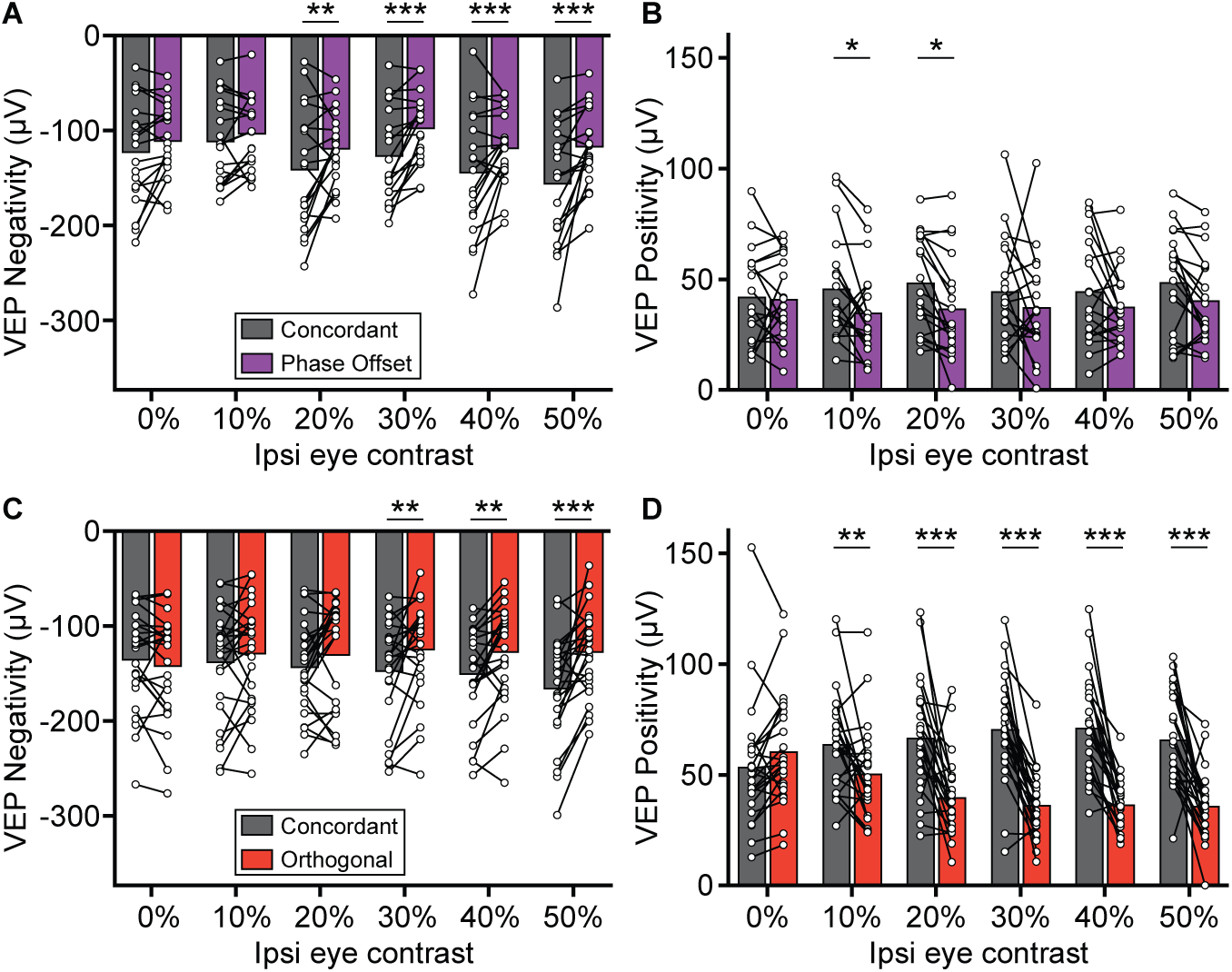
Phase offset and orthogonal stimuli affect different components of the VEP. **(A)** VEP negativities for Concordant (grey) and Phase Offset (purple) stimuli at different ipsilateral eye contrasts. Analyses performed using a two-way repeated measures ANOVA (main effect of condition, F(1,18) = 14.0, p = 0.002; condition × contrast, F(5,90) = 2.88, p = 0.019) followed by Šídák’s multiple comparisons test (concordant vs. phase offset at 0%, 10%, 20%, 30%, 40%, and 50% ipsi eye contrast: p = 0.393, 0.800, 0.009, <0.001, 0.001, <0.001). **(B)** VEP positivities for Concordant and Phase Offset stimuli at different ipsilateral eye contrasts. Two-way repeated measures ANOVA (main effect of condition, F(1,18) = 9.42, p = 0.007; condition × contrast, F(5,90) = 1.03, p = 0.407) followed by Šídák’s multiple comparisons test (concordant vs. phase offset at 0%, 10%, 20%, 30%, 40%, and 50% ipsi eye contrast: p = >0.999, 0.029, 0.015, 0.304, 0.349, 0.176). **(C)** VEP negativities for Concordant (grey) and Orthogonal (red) stimuli. Two-way repeated measures ANOVA (main effect of condition, F(1,23) = 11.8, p = 0.002; condition × contrast, F(5,115) = 6.09, p < 0.001) followed by Šídák’s multiple comparisons test (concordant vs. discordant at 0%, 10%, 20%, 30%, 40%, and 50% ipsi eye contrast: p = 0.847, 0.586, 0.210, 0.002, 0.002, <0.001). **(D)** VEP positivities for Concordant and Orthogonal stimuli. Two-way repeated measures ANOVA (main effect of condition, F(1,23) = 48.73, p < 0.001; condition × contrast, F(5,115) = 15.4, p < 0.001) followed by Šídák’s multiple comparisons test (concordant vs. discordant at 0%, 10%, 20%, 30%, 40%, and 50% ipsi eye contrast: p = 0.437, 0.009, <0.001, <0.001, <0.001, <0.001).

Because we saw more robust reductions in the VEP for orthogonal stimulus orientations than for phase offset disparities, we next sought to further characterize the features of VEP responses elicited by this stimulus combination. First, we examined how sensitive this effect was to graded orientation disparities between the eyes. Fixing the contralateral eye grating stimulus at 100% contrast at a 45° orientation and the ipsilateral eye contrast at 50%, we varied the grating orientation of the ipsilateral eye stimulus in 5° increments (from 0° to 30°, plus a 90° condition relative to the contralateral eye) (**Figure 3A**). Interocular orientation differences of as little at 10° showed a statistically significant decrease relative to the 0° (concordant) condition (**Figure 3B**), indicating that the VEP is highly sensitive to interocular orientation disparity.

**Figure 3.**
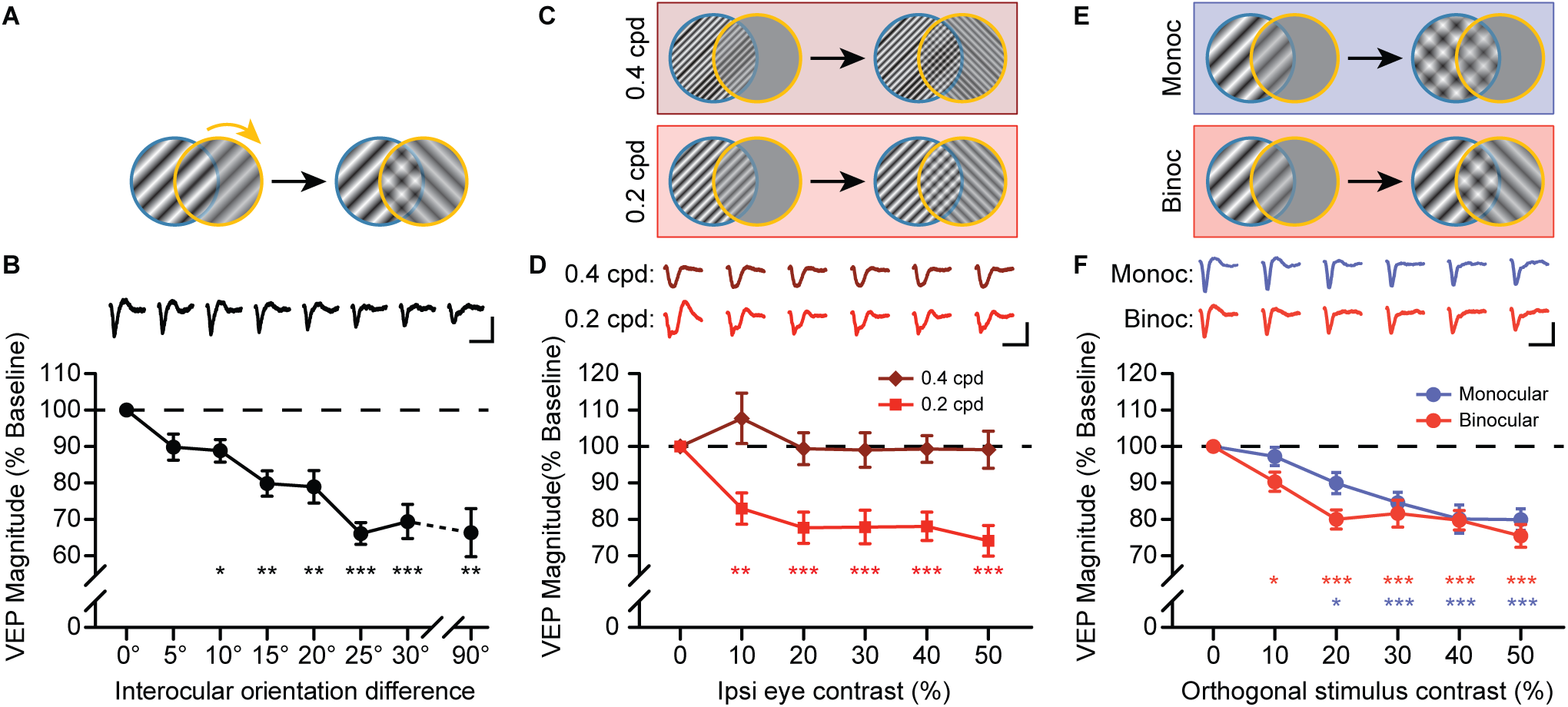
Reductions in VEP magnitude for discordant stimuli are sensitive interocular orientation differences, spatial frequency, and binocularity. **(A)** Mice viewed dichoptic grating stimuli (100% contra and 50% ipsi eye contrast) with varying interocular orientation differences (0° to 30° in 5° increments, plus an orthogonal orientation). VEP magnitudes were normalized to the baseline (0°) condition. Orientation differences of 10° or larger elicited significantly smaller VEPs. One-way repeated measures ANOVA with Geisser-Greenhouse correction (F(3.32, 29.8) = 12.7, p < 0.001) followed by Dunnett’s multiple comparisons test (0° vs. 5°, 10°, 15°, 20°, 25°, 30°, and 90°: p = 0.086, 0.025, 0.001, 0.005, <0.001, <0.001, 0.003). **(C)** Mice viewed discordant stimuli at higher spatial frequencies (0.2 and 0.4 cpd) with orthogonal ipsi eye stimuli (100% contra eye contrast, 0% to 50% ipsi eye contrast). **(D)** VEP magnitude was reduced for 0.2 cpd, but not for 0.4 cpd. Two-way repeated measures ANOVA with Geisser-Greenhouse correction (main effect of SF, F(1,20) = 22.66, p < 0.001; SF × contrast, F(2.92,58.4) = 4.69, p = 0.006) followed by Dunnett’s multiple comparisons test (0% vs. 10%, 20%, 30%, 40%, and 50% ipsi eye contrast for 0.02 cpd: p = 0.003, <0.001, <0.001, <0.001, <0.001; for 0.04 cpd: p = 0.701, >0.999, >0.999, >0.999, >0.999). **(E)** Mice viewed dichoptic grating stimuli (100% contra eye contrast) with a conflicting orthogonal grating. The orthogonal grating (0% to 50% contrast) was either presented to the ipsi eye (Binocular) or overlayed on top of the initial stimulus orientation for the contra eye (Monocular). **(F)** VEP magnitude for Binocular and Monocular conditions decreased as orthogonal grating contrast increased, but this decrease was sensitive to lower stimulus contrast in the Binocular condition. Two-way repeated measures ANOVA with Geisser-Greenhouse correction (main effect of condition, F(1,23) = 4.38, p = 0.048; condition × contrast, F(3.48, 80.0) = 1.44, p = 0.235) followed by Dunnett’s multiple comparisons test (0% vs. 10%, 20%, 30%, 40%, and 50% contrast for Monocular: p = 0.817, 0.011, <0.001, <0.001, <0.001; for Binocular: p = 0.006, <0.001, <0.001, <0.001, <0.001). **B,D,F:** Error bars indicate SEM. Scale bars: 200 ms and 100 µV.

Next, we examined the orientation disparity effect at higher spatial frequencies. We again presented orthogonal stimulus orientations to the eyes, with the contralateral eye fixed at 100% contrast, and the ipsilateral eye ranging from 0% to 50% contrast. Stimuli were presented at either 0.2 or 0.4 cycles per degree (cpd) (**Figure 3C**). At 0.2 cpd we again observed a strong reduction in VEP magnitude relative to the 0% contrast baseline, but no change was observed for 0.4 cpd (**Figure 3D**). This suggests that rivalrous stimulus effects on the VEP are spatial frequency-dependent.

Finally, we asked whether the reduction in VEP magnitude observed for orthogonal binocular stimuli could be accounted for by interocular interactions alone, or whether similar effects might also be elicited by overlapping stimuli within a single eye. To test this, we presented a full contrast grating to the contralateral eye, and added an overlapping orthogonal grating to either the same eye (monocular condition) or to the opposite eye (binocular condition), with contrast ranging from 0% to 50% (**Figure 3E**). In both cases, the superimposed gratings created a plaid-like pattern, differing only in whether the component gratings were presented to one or both eyes. While VEP magnitude decreased with increasing contrast in both conditions, the reduction was more sensitive to contrast in the binocular condition (**Figure 3F**). These results suggest that interocular presentation contributes additional suppression beyond that observed for monocular cross-oriented stimuli, consistent with a role for binocular interactions in shaping V1 responses to discordant inputs.

### Changes in the VEP are driven by an early decrease in V1 firing with phase disparity, and prolongation of firing with orientation disparity

While VEPs are a useful method for measuring summed neuronal responses within V1, one cannot infer what circuitry underpinnings may explain differential, stimulus-dependent VEP waveform changes. To better answer this question, we used 64-channel laminar probes to measure neuronal activity across all layers of V1 (**Figure 4A**). Mice viewed dichoptic phase reversing grating stimuli as described previously (see **Figure 1B**). The eye contralateral to the recording electrode viewed full contrast stimuli at a fixed orientation, while the ipsilateral eye viewed grey screen (monocular condition), an in-phase grating stimulus at the same angle (concordant condition), an out-of-phase grating stimulus at the same angle (phase offset condition), or an orthogonal grating (orthogonal condition, **Figure 4B**). In order to test all conditions within each animal, ipsilateral eye stimuli were presented only at 50% contrast for each dichoptic condition.

**Figure 4.**
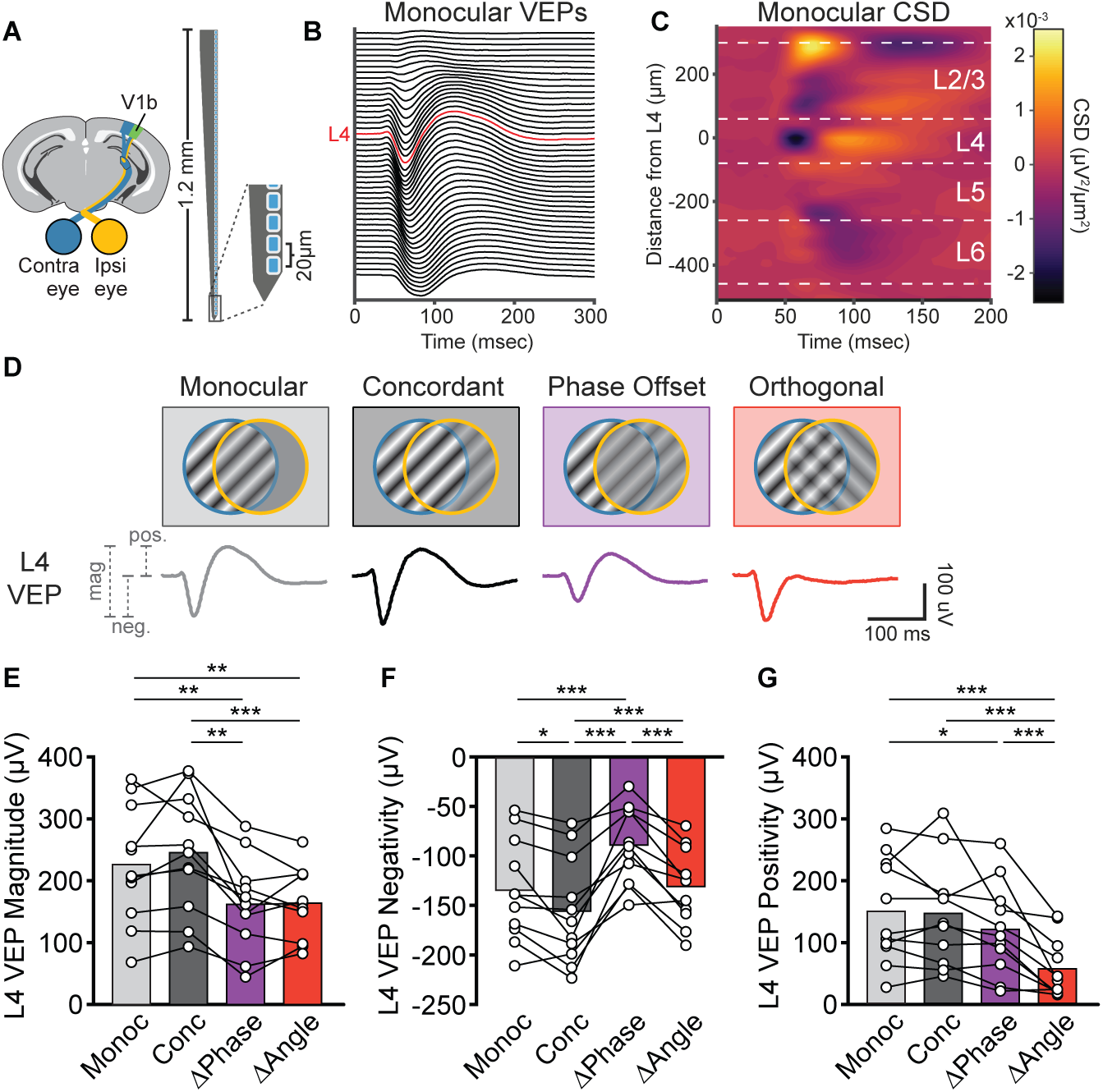
L4 VEPs measured using laminar probes reveal reductions in different components of the VEP for phase offset and discordant stimuli. **(A)** Recording setup. LFPs were recorded from bV1 using a 64-channel silicon probe. The probe spanned all cortical layers with channel spacing of 20 µm. **(B-C)** Monocular VEPs recorded from each electrode channel (**B**) were used to calculate current source densities (CSDs) shown in (**C**). The earliest current sink in the CSD for each mouse was used to identify the electrode corresponding to layer 4 (shown in red in **B**). Approximate layer boundaries shown as white dashed lines in **C**. **(D)** Stimulus conditions. Mice viewed dichoptic phase reversing gratings (2 Hz, 0.05 cpd) as described previously. Contra eye stimuli (45°, 100% contrast) were kept constant across conditions. Ipsi eye stimuli varied across 4 conditions: Monocular (0% contrast), Concordant (50% contrast, 45°), Phase Offset (50% contrast, 45° with 180° phase offset), and Orthogonal (50% contrast, 135°). L4 VEPs for each condition shown below schematic. **(E)** VEP magnitude for the Phase Offset (ΔPhase) and Orthogonal (ΔAngle) conditions were decreased relative to the Monocular (Monoc) and Concordant (Conc) conditions. One-way repeated measures ANOVA with Geisser-Greenhouse correction (F(2.20, 22.0) = 20.0, p < 0.001) followed by Tukey’s multiple comparison’s test. **(F)** Relative to Monocular, the negative component of the L4 VEP was larger (more negative) for Concordant, smaller for Phase Offset, and no different for Orthogonal stimuli. One-way repeated measures ANOVA with Geisser-Greenhouse correction (F(2.15, 21.5) = 24.9, p < 0.001) followed by Tukey’s multiple comparison’s test. **(G)** Relative to monocular, the positive peak of the layer 4 VEP was slightly decreased for Phase Offset stimuli, and substantially diminished for Orthogonal stimuli. One-way repeated measures ANOVA with Geisser-Greenhouse correction (F(2.16, 21.6) = 19.7, p < 0.001) followed by Tukey’s multiple comparison’s test. **E-G:** *p<0.05, **p<0.01, ***p<0.001.

Recording VEPs across all channels enabled us to calculate a current source density plot for each mouse. The electrode contact corresponding to L4 was identified for each mouse based on the earliest current sink in the current source density (CSD) plots (**Figure 4C**, see Methods). Consistent with our results using unipolar Tungsten electrodes (**Figure 1**), we observed a reduction in L4 VEP magnitude for the phase offset and orthogonal conditions relative to concordant stimuli (**Figure 4D-E**). We also found a statistically significant decrease relative to monocular stimuli, which was previously only observed for orthogonal stimuli. Again, the reduction in VEP magnitude in the phase offset and discordant conditions were caused by distinct changes in VEP waveform; phase offset stimuli primarily decreased the VEP negativity, whereas discordant stimuli primarily decreased the VEP positivity (**Figure 4F-G**).

To determine how changes in different components of the VEP relate to changes in neuronal spiking in bV1, we next analyzed unit activity across cortical layers. Recordings from individual mice were aligned based on the channel that showed the earliest L4 current sink, and the multiunit activity envelope (MUAe) was calculated for each electrode contact (see Methods). Approximate layer boundaries were assigned based on the distance from the L4 sink and the L5 peak in MUAe (Senzai, Fernandez-Ruiz et al. 2019) (**Figure 5A**). To account for the variability in spiking between layers, we z-scored the MUAe for each channel based on its average across the entire recording session. We observed a strong visually evoked response across all layers of V1 when mice viewed the grating monocularly through the contralateral eye (**Figure 5B**). This response peaked around 60-80 ms after stimulus presentation and MUAe returned to near-baseline levels by ∼100 ms. To compare how activity across layers was affected by the different stimulus conditions, we then assessed MUAe when the ipsilateral eye viewed a concordant, phase offset, or orthogonal grating stimulus (**Figure 5C**). MUAe changed across most layers of V1 for all three conditions (**Figure 5D**). Relative to monocular stimuli, concordant and orthogonal stimuli increased the response in the early period following the phase reversal (roughly corresponding in time with the negative component of the VEP), whereas phase offset stimuli elicited a reduced response in this period (**Figure 5E-F**). Unexpectedly, the orthogonal condition prolonged firing well beyond the visually evoked response of the other conditions (**Figure 5E,G**). This prolonged period of firing persisted through 200 ms after the phase reversal, corresponding to the timing of the VEP positivity in response monocular and concordant stimuli (**Fig 4**). Thus, the reduction in the positive component of the VEP waveform by the orthogonal condition may reflect prolongation of activity.

**Figure 5.**
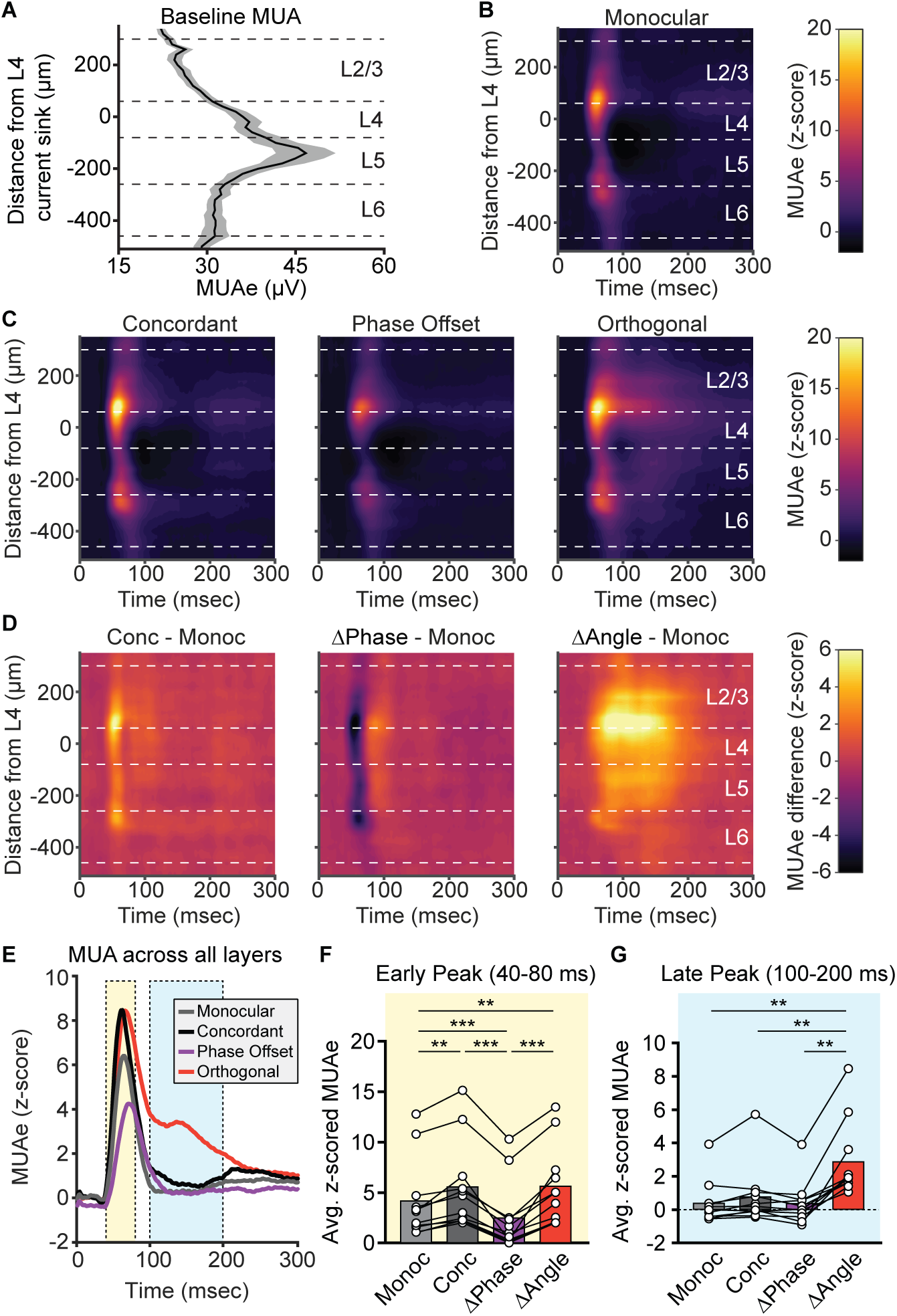
Multiunit activity across layers of V1 reveals changes in early and late firing that differs for phase offset and orthogonal stimuli. Multiunit activity was measured across layers of bV1 during the experiment shown in Figure 3. **(A)** Plot of the multiunit activity envelope (MUAe) for each channel during a baseline period. MUAe was greatest in L5. **(B-C)** MUAe for each condition was z-scored based on the baseline MUAe shown in **A**. The z-scored MUAe for all conditions showed a brief short-latency peak in activity corresponding in time to the VEP negativity. All cortical layers showed a statistically meaningful visual response (z-score > 1.96). **(D)** The z-scored MUAe for each of the conditions in **C** were compared with the Monocular (Monoc) condition in **B**. Concordant (Conc) stimuli elicited a larger early peak in activity across cortical layers, Phase Offset (ΔPhase) stimuli elicited a reduced early activity across all layers, and Orthogonal (ΔAngle) stimuli elicited an extended period of elevated activity, particularly in L2/3-L5. **(E)** Due to the similarity in the changes in activity observed across layers of V1, we collapsed the z-scored MUAe across all layers to directly compare differences in activity in the early (40-80 ms, yellow) and late (100-200 ms, blue) periods for each condition. **(F)** In the early period following each phase reversal, Concordant and Orthogonal stimuli elicited greater activity than the Monocular condition, and Phase Offset stimuli elicited reduced activity. One-way repeated measures ANOVA with Geisser-Greenhouse correction (F(1.54, 15.4) = 37.7, p < 0.001) followed by Tukey’s multiple comparisons test. **(G)** In the late period following the initial peak in activity, Orthogonal stimuli elicited increased activity relative to each of the other conditions. One-way repeated measures ANOVA with Geisser-Greenhouse correction (F(1.28, 12.8) = 25.4, p < 0.001) followed by Tukey’s multiple comparisons test. Dashed lines in **A-D** indicate approximate borders between cortical layers. **F** and **G**: **p<0.01, ***p<0.001.

### Both regular-spiking and fast-spiking units show prolonged activity with orthogonal stimuli

Based on our finding that dichoptic orthogonal grating stimuli elicit prolonged MUA responses, we hypothesized that this persistent activity could result in a reduction in inhibitory drive. Specifically, we reasoned that if inhibitory interneuronal activity was diminished during this period, excitatory neurons could remain active for a longer period. To explore this possibility, we first examined parvalbumin-positive (PV+) interneurons, the most abundant inhibitory cell type in cortex and key mediators of fast, perisomatic inhibition onto pyramidal neurons (Tremblay, Lee et al. 2016). Although PV+ interneurons often exhibit similar stimulus tuning to pyramidal neurons (Niell and Stryker 2008), we hypothesized that this relationship might be disrupted during binocular conflict. If disinhibition contributes to the prolonged firing observed for orthogonal stimuli, it might be reflected in altered PV+ cell activity during this window. To test this, we classified single units as fast-spiking (FS, putative PV+ interneurons) or regular-spiking (RS, putative pyramidal) based on trough-to-peak latency (**Figure 6A**), and compared their responses across conditions (**Figure 6B**). RS units exhibited a clear early peak in firing (40-80 ms after stimulus onset) that was elevated for concordant and orthogonal stimuli, and reduced for phase offset stimuli (**Figure 6C-D**). Similar to the MUA results, a prolonged increase in RS activity during the 100-200 ms window was observed only for orthogonal stimuli (**Figure 6C, E**). However, FS units showed a strikingly similar pattern of responses (**Figure 6F-H**), suggesting that PV+ interneurons are not selectively disengaged during the late phase of orthogonal stimulation. Given this parallel between putative excitatory and PV+ cell activity, we next considered whether the structure of these responses varied by cortical layer, which could offer further insight into how binocular conflict is processed across the bV1 circuit.

**Figure 6.**
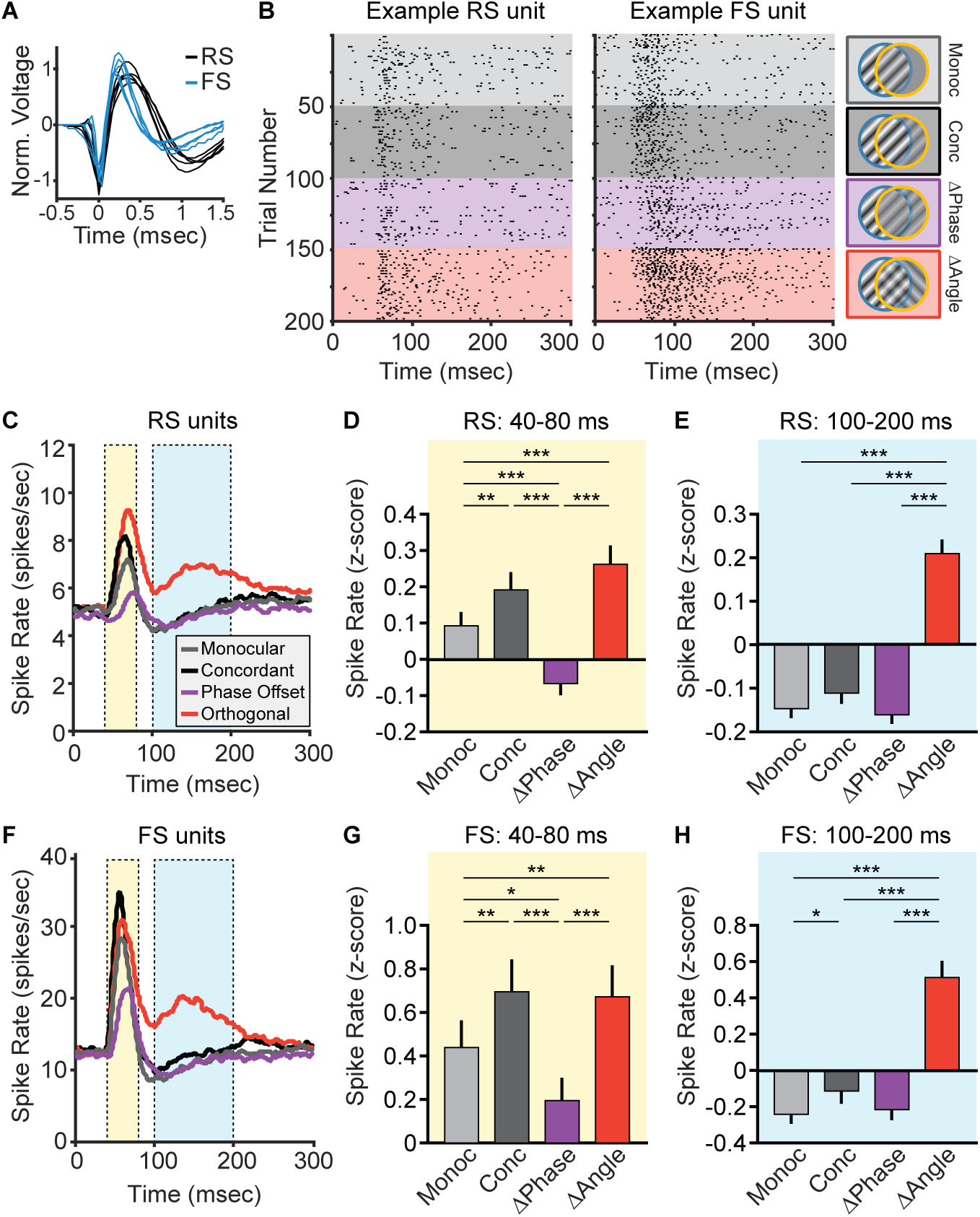
Regular and fast spiking units in V1 show similar responses across conditions. **(A)** Single units were classified as regular spiking (RS, black) or fast spiking (FS, blue) based on their trough-to-peak width (see Methods). **(B)** Raster plots of action potentials for a representative RS unit and an example FS unit for a subset of trials during each of the 4 stimulus conditions: Monocular (Monoc, light grey), Concordant (Conc, dark grey), Phase Offset (ΔPhase, purple), and Orthogonal (ΔAngle, red). **(C)** PSTH for all RS units (putative principal cells, n=275) averaged across all stimuli in each condition. **(D)** Average firing rate for RS units during the early peak in activity (40-80 ms after stimulus presentation, yellow shaded region in **C**). Relative to the Monocular condition, Concordant and Orthogonal stimuli showed an increase in early RS activity, and Phase Offset stimuli showed decreased activity. One-way repeated measures ANOVA with Geisser-Greenhouse correction (F(2.03, 557) = 33.78, p < 0.001) followed by Tukey’s multiple comparisons test. **(E)** Average firing rate for RS units during the late period of activity (100-200 ms after stimulus presentation, blue shaded region in **C**). Orthogonal stimuli elicited increased firing relative to the other three conditions. One-way repeated measures ANOVA with Geisser-Greenhouse correction (F(1.88, 516) = 57.0, p < 0.001) followed by Tukey’s multiple comparisons test. **(F)** PSTH for all FS units (putative PV+ neurons, n=52) averaged across all stimuli in each condition. **(G)** Average firing rate of FS units 40-80 ms after stimulus presentation (yellow shaded region in **F**). Concordant and Orthogonal stimuli elicited increased activity and Phase Offset stimuli elicited decreased activity relative to the Monocular condition. One-way repeated measures ANOVA with Geisser-Greenhouse correction (F(1.99, 102) = 14.9, p < 0.001) followed by Tukey’s multiple comparisons test. **(H)** Average firing rate of FS units 100-200 ms after stimulus presentation (blue shaded region in **F**). Orthogonal stimuli elicited increased firing relative to all other conditions. One-way repeated measures ANOVA with Geisser-Greenhouse correction (F(1.59, 81.0) = 27.6, p < 0.001) followed by Tukey’s multiple comparisons test. **D,E,G,H:** *p<0.05, **p<0.01, ***p<0.001

To explore this, we analyzed the laminar distribution of RS and FS unit activity across conditions to determine whether stimulus-dependent differences in firing rates were uniformly expressed throughout cortex or varied by layer. Single units were sorted by depth and assigned to L2/3, L4, L5, and L6 based on their location relative to L4 (RS: **Figure S1**; FS: **Figure S2**). In the early time window (40–80 ms after stimulus onset), RS firing rates differed by both stimulus condition and cortical layer. In particular, responses to phase-offset stimuli were reduced in L2/3, L5, and L6 compared with other conditions, while responses in L4 were relatively unchanged (**Figure 7A**). This observation was confirmed by a significant condition-by-layer interaction. FS units, in contrast, showed no such interaction during the early period, with broadly similar responses across layers and conditions (**Figure 7B**). During the late time window (100–200 ms), RS units showed elevated firing for orthogonal stimuli across all layers (**Figure 7C**), consistent with the MUA results (**Figure 5B**). FS units also responded more strongly to orthogonal stimuli during this period, but this effect was not uniform across lamina. Instead, it was most pronounced in L2/3, where responses to orthogonal stimuli were significantly greater than those to other conditions (**Figure 7D**). Together, these findings show that suppression of RS activity by phase-offset stimuli occurs selectively outside of L4, suggesting that this effect is not inherited from thalamic input but instead emerges within intracortical circuitry. In contrast, the enhancement of RS and FS activity by orthogonal stimuli occurs broadly across layers for RS units but is selectively amplified in L2/3 for FS units, pointing to a potential layer- and cell-type– specific locus of circuit engagement during binocular orientation conflict.

**Figure 7.**
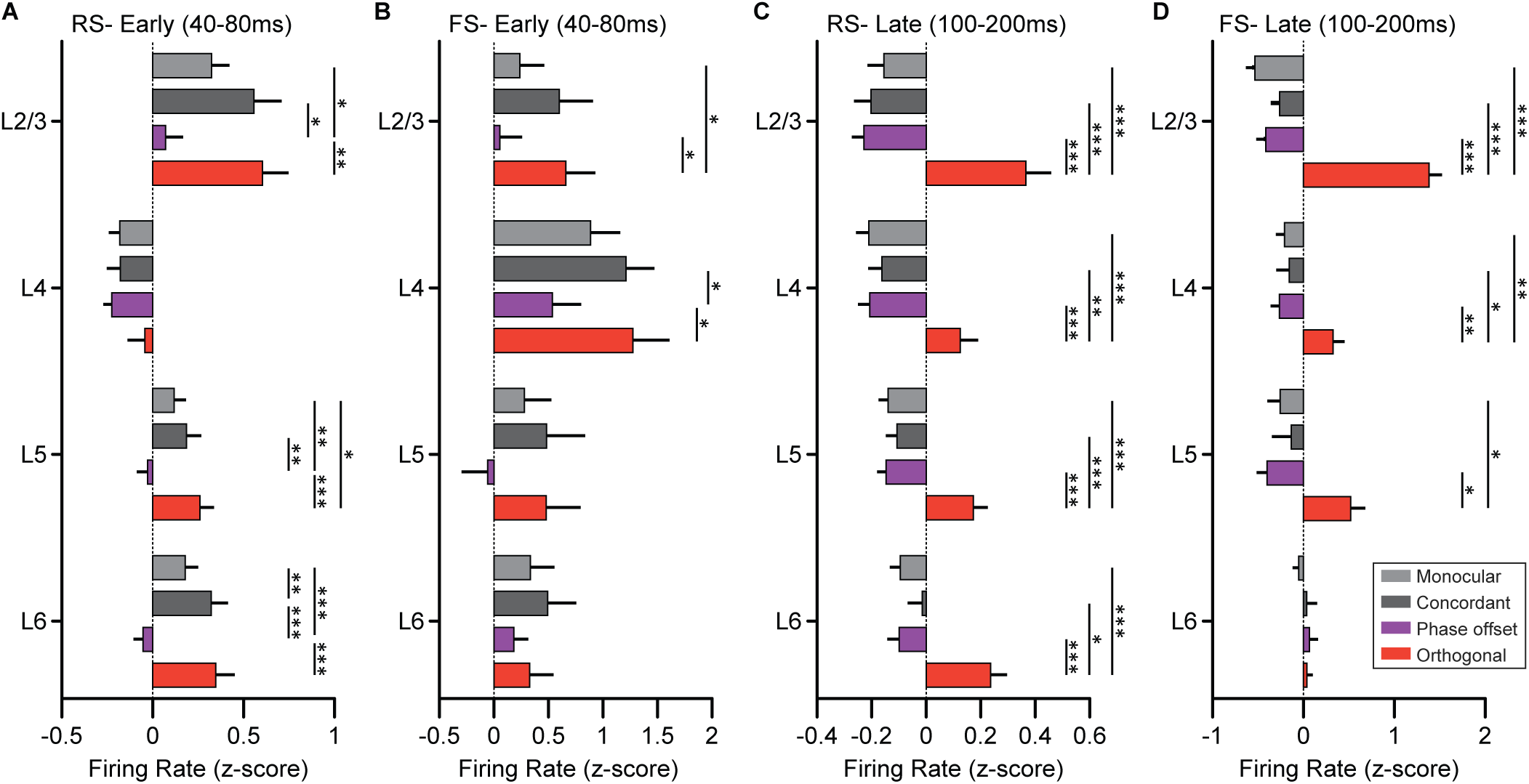
Laminar differences in responses of RS and FS units during early and late periods. **(A)** Mean firing rates of RS units in each cortical layer during the early visually evoked response (40–80 ms post-stimulus) under Monocular (light gray), Concordant (dark gray), Phase Offset (purple), and Orthogonal (red) conditions. A significant interaction between stimulus condition and layer was observed (two-way repeated measures ANOVA with Geisser-Greenhouse correction: main effect of condition, F(2.05, 557) = 36.9, p < 0.001; layer, F(3, 271) = 8.27, p < 0.001; condition × layer, F(9, 813) = 2.54, p = 0.007), driven by reduced responses to Phase Offset stimuli in L2/3, L5, and L6 but not in L4. **(B)** FS unit responses during the early time window. No significant interaction between condition and layer was found (main effect of condition, F(1.99, 95.4) = 15.9, p < 0.001; layer, F(3, 48) = 1.85, p = 0.15; condition × layer, F(9, 144) = 1.03, p = 0.41), indicating relatively uniform responses across layers. **(C)** RS unit firing rates during the late response window (100-200 ms). Firing rates were significantly modulated by condition (F(1.89, 513) = 60.7, p < 0.001), with elevated responses to Orthogonal stimuli observed across all layers. A main effect of layer was also observed (F(3, 271) = 2.66, p = 0.048), but there was no significant condition × layer interaction (F(9, 813) = 1.53, p = 0.13). **(D)** FS unit firing rates during the late time window. A significant condition-by-layer interaction was found (main effect of condition, F(2.14, 103) = 58.9, p < 0.001; layer, F(3, 48) = 0.64, p = 0.59; condition × layer, F(9, 144) = 14.5, p < 0.001), driven by a pronounced increase in L2/3 FS activity in response to Orthogonal stimuli. **A-D:** Error bars indicate SEM. Šídák’s multiple comparisons test used for post-hoc comparisons. *p<0.05, **p<0.01, ***p<0.001.

### bV1 principal cells and somatostatin-positive interneurons show opposite responses to orthogonal dichoptic stimuli

In the context of established cortical circuitry (Tremblay, Lee et al. 2016), the similarity in responses between RS and FS units to the different dichoptic stimulus conditions suggests that modulation of PV+ interneurons is unlikely to mediate the persistent activity triggered by the orthogonal condition. The next most likely candidate was somatostatin-positive (SOM+) interneurons, which inhibit both excitatory cells and PV+ interneurons in the mammalian cortex via feedback inhibition. We hypothesized that a decrease in SOM+ cell activity elicited by the orthogonal condition disinhibits both PV+ and principal cells (PCs) in bV1. To compare the responses of genetically defined PCs and SOM+ interneurons we used 2P calcium imaging. Mice expressing GCaMP in promoter-specific neurons were head-fixed in front of a dichoptic display viewing the same stimuli as described above (**Figure 8A,B**). Visual responses of individual neurons were measured using a 2P microscope (**Figure 8C**). First, we genetically expressed GCaMP6f in excitatory neurons using an EMX1-cre x GCaMP6f.tTA mouse line and measured the responses at a depth corresponding to L2/3 (**Figure 8D**). Due to the relatively slow temporal dynamics of GCaMP, we expected that the measured change in fluorescence (ΔF/F) response for each condition would reflect the overall differences in firing rate that we observed in the MUA, favoring a difference for the orthogonal condition. This is indeed what we observed. Although there was a substantial amount of variation in response to the grating stimulus orientation presented (**Figure 8E**), on average there was a robust increase in responses when the orthogonally oriented grating was presented to the ipsilateral eye and with each phase reversal (**Figure 8F-H**).

**Figure 8.**
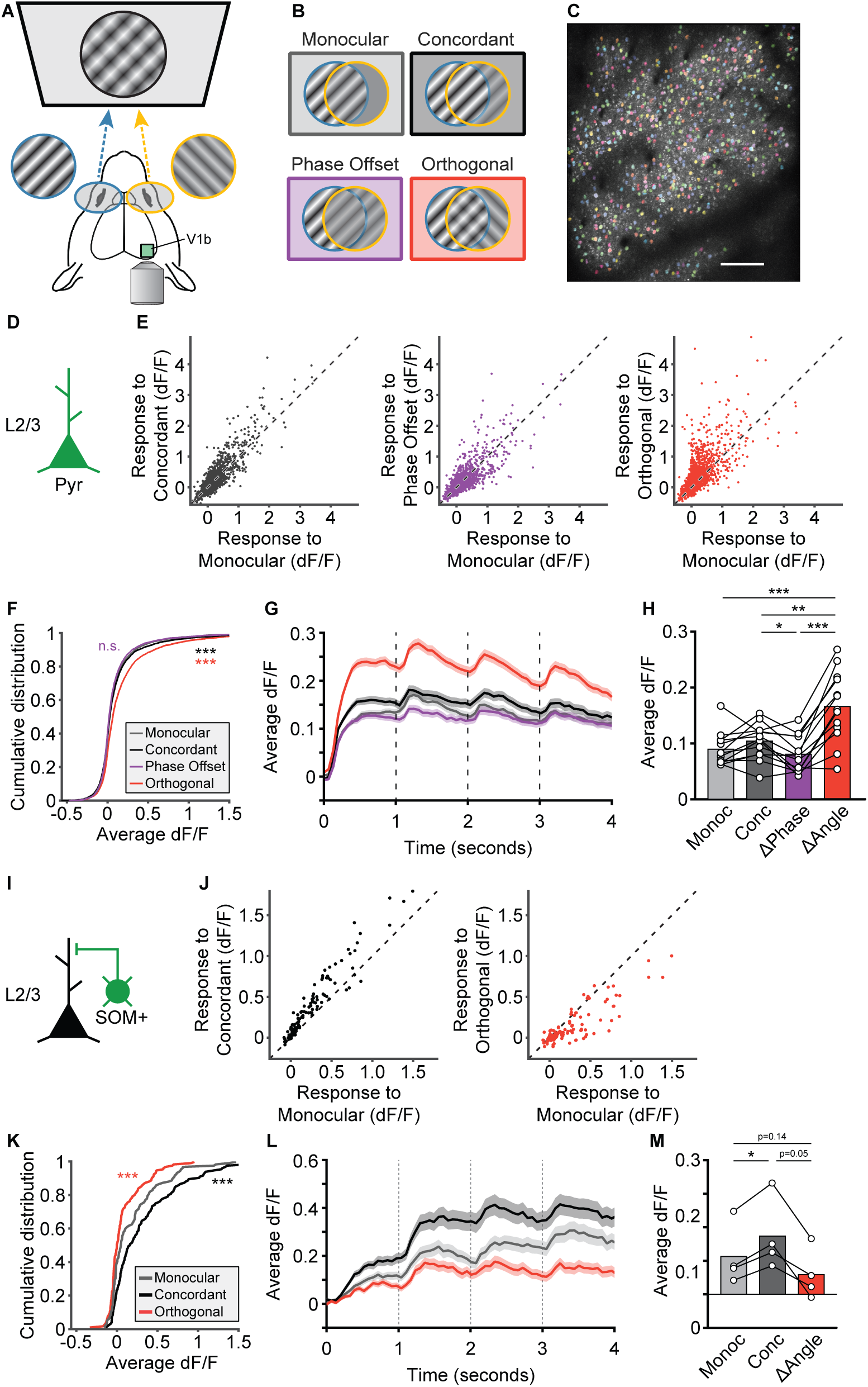
Orthogonal stimuli elicit larger calcium responses in L2/3 principal cells, but reduced calcium responses in SST+ interneurons. **(A)** Recording setup. Head-fixed mice expressing GCaMP with a cranial window above bV1 viewed phase-reversing grating stimuli on a dichoptic display. **(B)** Experimental conditions. Mice viewed Monocular (contra only), Concordant (contra and ipsi at same phase and orientation), Phase Offset (180° offset phase, same orientation), and Orthogonal (90° orientation difference) stimuli. **(C)** Example FOV for a mouse expressing GCaMP6f in excitatory (Emx1+) cells imaged 200µm below pia (L2/3). Colored patches indicate ROIs for individual neurons. Scale bar=100µm. **(D-H)** Responses of L2/3 pyramidal cells to each of the 4 conditions. **(E)** Scatter plots showing average responses of individual cells to Concordant (black), Phase Offset (purple), and Orthogonal (red) stimuli relative to Monocular. dF/F was calculated relative to baseline response during interblock intervals. Dashed identity line on each plot indicates an equal response to both conditions. **(F)** Cumulative distribution plot comparing the responses of all neurons across conditions. One-way repeated measures ANOVA with Geisser-Greenhouse correction (F(2.59, 9724) = 124.2, p<0.001) followed by Tukey’s multiple comparisons test (***p<0.001 relative to Monocular). **(G)** Plot showing population response to each stimulus across all blocks for each condition. Shaded error bars indicate SEM. **(H)** Comparison of responses for each condition by mouse (n=13). One-way repeated measures ANOVA with Geisser-Greenhouse correction (F(1.86, 22.28) = 27.2, p < 0.001) followed by Tukey’s multiple comparisons test (*p<0.05, **p<0.01, ***p<0.001). **(I-M)** Responses of L2/3 SOM+ interneurons expressing GCaMP7f for Monocular, Concordant, and Orthogonal stimulus conditions. **(J)** Scatter plots showing average responses of individual cells to Concordant (black) and Orthogonal (red) stimuli relative to Monocular. Dashed identity line on each plot indicates an equal response to both conditions. **(K)** Cumulative distribution plot comparing the responses of all SOM+ cells across conditions. One-way repeated measures ANOVA with Geisser-Greenhouse correction (F(1.23, 166.5) = 95.5, p<0.001) followed by Tukey’s multiple comparisons test (***p<0.001 relative to Monocular). **(L)** Plot showing response of all SOM+ cells across time for each stimulus condition. Shaded error bars indicate SEM. **(M)** Comparison of responses for each condition by mouse (n=4). One-way repeated measures ANOVA with Geisser-Greenhouse correction (F(1.03, 3.11) = 15.7, p=0.026) followed by Tukey’s multiple comparisons test (*p<0.05).

Next, we tested whether SOM+ cells might be differentially engaged by orthogonal dichoptic stimuli. We expressed GCaMP7f in SOM+ interneurons in bV1 by injecting an AAV expressing cre-dependent GCaMP7f into mice on a SOM-cre genetic line and imaged SOM+ cell ΔF/F in L2/3 (**Figure 8I**). This time, we only presented mice with the monocular, concordant, and orthogonal conditions and observed a robust and uniform increase in responses for concordant stimuli relative to monocular stimuli and a decrease in responses for orthogonal stimuli that grew with each phase reversal (**Figure 8J-M**). These observations draw a clear contrast between the responses of genetically defined neuronal subtypes to reveal insights into the circuitry underling bV1 responses to rivalrous dichoptic stimuli.

## DISCUSSION

### Cortical responses to concordant and phase offset stimuli can be accounted for via feedforward mechanisms

In this study, we used a combination of VEPs (**Figures 1-4**), MUA (**Figure 5**), SUA (**Figures 6-7**), and 2P calcium imaging (**Figure 8**) to investigate how population cortical responses of mouse bV1 are modulated by different combinations of dichoptic visual stimuli. We found that relative to monocular stimuli, concordant binocular stimuli and 180° phase offset stimuli elicit opposing changes in the visually evoked response. Concordant stimuli enhanced the early negative component of the VEP via a short latency increase in MUA, whereas phase offset stimuli reduced the VEP negativity through a corresponding decrease in MUA. This elevated response to concordant stimuli likely reflects increased drive of binocular neurons due to the addition of inputs from the ipsilateral eye or the recruitment of the few ipsilateral-referring neurons within bV1 (**Figure 9**). The relatively modest increase in VEP and MUA responses for concordant relative to monocular stimuli likely reflects sublinear binocular integration in mouse V1, consistent with prior work demonstrating that binocular inputs do not sum linearly but are shaped by normalization mechanisms that preserve tuning and constrain response amplitude (Longordo, To et al. 2013, Zhao, Liu et al. 2013).

**Figure 9.**
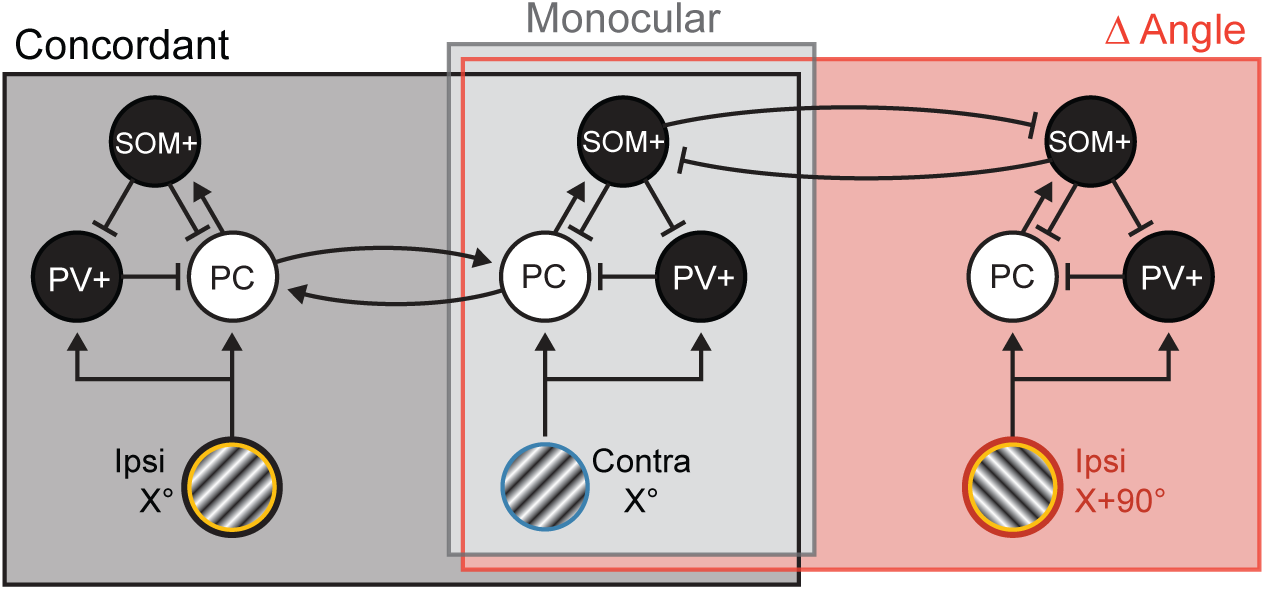
Circuit model for concordant and orthogonal responses. Proposed circuit diagram to account for responses elicited by concordant and conflicting visual stimuli. Inputs from the contra eye (light grey box) drive excitatory principal cells (PCs) and inhibitory PV+ interneurons. Feedback inhibition from SOM+ interneurons will constrain this response in time. Concordant inputs to the ipsi eye (dark grey box) will recruit a separate population of ipsi-responsive monocular PCs and PV+ cells. Ipsi and contra responsive cells are reciprocally connected. Ipsi eye inputs with an orthogonal orientation (red box) will also drive ipsi-responsive PCs and PV+ cells tuned to that orientation, but here there are not reciprocal connections between PCs. Instead, lateral inhibition between populations of SOM+ cells will reduce feedback inhibition onto PCs and PV+ cells, prolonging the evoked activity of these neurons.

The reduced response for phase offset stimuli can also be accounted for by known feedforward mechanisms. Binocular simple cells in mouse V1 receive spatially aligned inputs from both eyes that are typically tuned to the same orientation but differ in phase sensitivity. When gratings of identical orientation but opposite phase are presented dichoptically, these inputs can cancel due to destructive interference at the level of subthreshold membrane potential, as observed in previous whole-cell recordings (Zhao, Liu et al. 2013). This interaction reduces the net depolarization and consequently suppresses spiking output. Moreover, the amplitude-dependent sublinear integration described by Longordo et al. (2013) predicts that strong but conflicting monocular inputs, such as those generated by out-of-phase gratings, will not sum constructively and may even inhibit one another, further diminishing the cortical response. Together, these feedforward properties of binocular integration constrain responses to phase-disparate stimuli and likely contribute to the observed suppression in MUA and VEP amplitude.

This suppressive effect can also be interpreted in the context of spatial disparity. A 180° phase offset in grating stimuli at 0.05 cpd, as used in our experiments, results in a spatial disparity of 10° of visual field. The suppression of early visually evoked responses under this condition suggests a feed-forward cortical mechanism, consistent with the sublinear integration and cancellation described above. Importantly, this pattern may also reflect the disparity tuning properties of individual neurons in mouse bV1. Prior studies have shown that mice exhibit broader disparity tuning than other mammals, such as cats and primates, likely reflecting adaptations for near-field visual processing in species that interact with objects at close range (Scholl, Burge et al. 2013). Indeed, individual bV1 neurons in mice can show preference for disparities of 10° or more (Samonds, Choi et al. 2019), making it plausible that the 180° phase offset grating stimulus drives a small subpopulation of neurons but V1 on average is activated less effectively as compared with zero-disparity (concordant) stimuli.

These observations are consistent with the disparity energy model first described by Ohzawa, DeAngelis, and Freeman (Ohzawa, DeAngelis et al. 1990), in which monocular receptive fields with a slight spatial offset converge to form binocular receptive fields tuned to specific spatial or phase disparities. In this framework, responses are enhanced when the binocular input matches a neuron’s preferred disparity and attenuated when the disparity deviates from that preferred offset, due to partial cancellation of left and right eye signals via linear summation. Disparity tuning of neurons in mouse V1 has been demonstrated using both single-unit recordings and calcium imaging, along with a corresponding pattern of facilitation and attenuation for preferred and non-preferred disparities, respectively (Scholl, Burge et al. 2013, Scholl, Pattadkal et al. 2017, La Chioma, Bonhoeffer et al. 2020). Our results showing an enhanced population response for concordant binocular stimuli relative to monocular stimuli, and a suppressed response for 180° phase offset stimuli, are consistent with a greater proportion of neurons tuned near 0° disparity and fewer tuned near 180°, in line with previous findings that most mouse V1 neurons prefer in-phase stimuli (Fu, Tanabe et al. 2023). While previous studies have characterized sublinear summation at the level of individual neurons (Longordo, To et al. 2013, Zhao, Liu et al. 2013), our findings extend these principles to population-scale measures and demonstrate how disparity-dependent facilitation and suppression shape early cortical responses to dichoptic stimuli.

It is notable that the pattern of suppression of cortical responses for phase offset stimuli is only statistically significant outside of L4 of bV1 for RS units (**Figure 7A**). Mice differ from other mammals in that binocular inputs converge within thalamorecipient L4 in bV1 (Gordon and Stryker 1996). Our findings therefore suggest that disparity tuning may emerge after this early stage of binocular processing. However, because we are measuring population-level responses for each cortical layer we cannot rule out the possibility that a subset of neurons within dLGN or L4 might display disparity tuning that drives the responses at subsequent stages of processing within bV1. Further work will be required to fully understand how the cortical circuitry of the early visual pathway gives rise to disparity tuning to support the processing of stimuli with interocular phase differences.

### Disinhibition of V1 responses to orthogonal binocular stimuli is mediated by SOM+ interneurons

Unlike the responses to concordant and phase offset stimuli, the pattern of activity we observe for binocular stimuli with conflicting stimulus orientations cannot be accounted for by a simple feedforward model. In particular, we see a period of heightened MUA and SUA after the initial visually evoked response (100-200 ms after stimulus presentation, **Figure 6**) which corresponds to a flattening of the VEP positivity (**Figures 2, 4**). This increase is apparent across cortical layers, and in the responses of both RS (mostly excitatory) and FS (mostly PV+ inhibitory) units within bV1 (**Figure 7B, C**).

Prior studies have shown that binocular integration in mouse V1 is sublinear when the two eyes receive matched inputs, particularly at the preferred orientation, and that this mechanism helps preserve orientation selectivity (Longordo, To et al. 2013, Zhao, Liu et al. 2013). While those studies did not address interocular orientation conflict, they suggest that binocular responses should remain modest when summed across eyes. Our findings are consistent with this prediction during the early response phase, but diverge in the later time window, where we observe a robust and selective enhancement of activity for orthogonal stimuli. This pattern suggests a shift from sublinear integration to active disinhibition.

SOM+ interneurons, which inhibit both PCs and PV+ interneurons within cortex and are a powerful source of feedback inhibition (Tremblay, Lee et al. 2016, Del Rosario, Coletta et al. 2024), are well-situated to mediate this disinhibition. Our 2P calcium imaging results showed that SOM+ cell activity increases for concordant binocular stimuli and decreases for stimuli with orthogonal orientations (**Figure 8**). Based on this, we propose a model (**Figure 9**) in which SOM+ cells play a key role in shaping cortical responses to binocular stimuli with concordant or orthogonal orientations. In this model, monocular inputs to V1 drive both PCs and PV+ interneurons; the PCs then activate SOM+ interneurons, which in turn provide feedback inhibition to PCs and PV+ cells. This feedback inhibition keeps the visually evoked response constrained in time and would generate the similar pattern of responses we observe for RS and FS units. Presenting a concordant stimulus to the other eye would increase the drive onto PCs (and PV+ cells), and therefore elicit stronger SOM+ cell activity (**Figure 8J-M**) and feedback inhibition. In the case of orthogonal stimulus presentation, however, there is a reduction in the activation of SOM+ cells, leading to disinhibition of PCs and PV+ cells after the initial peak of activity. This disinhibition could arise from local connections between competing and mutually inhibitory populations of SOM+ cells within V1, or via feedback from other cortical areas, possibly mediated by vasoactive intestinal peptide-expressing interneurons. Further experiments will be required to disentangle these possibilities.

In humans, presenting discordant stimuli to each eye results in a repeated switching of perception of each stimulus, a phenomenon known as binocular rivalry (Levelt 1965). Recordings of V1 during binocular rivalry experiments reveal modulation of activity corresponding to the perceived stimulus (Polonsky, Blake et al. 2000, Tong and Engel 2001). In awake monkeys presented with orthogonal grating stimuli to each eye and trained to report the perceived orientation, many neurons within V1 with a preference for one orientation respond only when the monkey reports perceiving that orientation, and are suppressed when the opposing percept is dominant (Leopold and Logothetis 1996). Presenting rivalrous stimuli to anesthetized monkeys elicits an alternating activation and suppression of ocular dominance columns corresponding to each eye that resembles the dynamics and pattern of switching that occurs in binocular rivalry, suggesting that rivalry within V1 is unlikely to solely rely on top-down mechanisms (Xu, Han et al. 2016). In contrast, V1 neurons recorded from strabismic monkeys did not show any net facilitation or suppression (Economides, Adams et al. 2021), suggesting that interocular suppression in strabismus is likely to be mediated at a higher level of processing.

Whether binocular rivalry occurs in mice, either at the level of perception or cortical dynamics, remains an open question. Our findings show that discordant binocular inputs elicit divergent responses in PV+ and SOM+ interneurons, with increased PV+ cell firing and suppressed SOM+ cell activity relative to concordant stimuli. These cell-type-specific dynamics suggest that mouse V1 circuits respond selectively to interocular conflict and may contribute to early-stage processing of binocular mismatch. While not a direct readout of perceptual suppression, these responses highlight fundamental principles of how cortical circuits handle interocular conflict. Given that binocular rivalry in humans is modulated by GABAergic signaling (Mentch, Spiegel et al. 2019), these findings underscore the value of the mouse as an accessible model system for dissecting the inhibitory circuit mechanisms that govern binocular integration and competition.

### Integration across modalities and implications for binocular processing

Although VEPs, unit recordings, and calcium imaging each capture different aspects of cortical activity, our findings highlight how these modalities together can clarify the temporal and cell-type-specific mechanisms underlying binocular processing. VEPs primarily capture the net synaptic currents across large populations of neurons and are particularly sensitive to changes in activity over time, rather than absolute firing rates (Hayden, Finnie et al. 2023). In this context, a reduction in VEP amplitude can arise from different underlying patterns. For example, a lower initial peak in firing can reduce both the early negative and subsequent positive components of the VEP, while a delayed or sustained increase in firing may decrease the relative change in activity over time, resulting in a flattened or diminished VEP positivity.

This framework helps reconcile several of our findings. For phase-offset stimuli, the reduction in VEP negativity aligns with decreased early MUA and SUA, suggesting a lower peak response. In contrast, orthogonal stimuli produce a reduced VEP positivity alongside an increase in late-phase spiking, reflecting a prolonged or redistributed firing pattern that diminishes the temporal contrast captured by the VEP. Calcium imaging of excitatory neurons supports this interpretation. Because calcium signals reflect integrated activity over longer timescales, the observed increase in fluorescence in response to orthogonal stimuli is consistent with the elevated firing rates during both early and late response windows. This result aligns with prior findings (Kim, Chaloner et al. 2020) that VEP magnitude does not simply reflect overall excitatory activity as measured by calcium imaging but is instead shaped by the timing and synchrony of neuronal responses.

These modality-specific differences underscore the importance of a multimodal approach for dissecting the dynamics of binocular integration. Our results show that different forms of interocular conflict engage distinct circuit mechanisms within mouse V1. Phase-offset stimuli rely on feedforward integration consistent with disparity tuning, whereas orientation conflicts recruit mechanisms involving specific cell types and cortical layers, consistent with disinhibition. These findings reveal how local circuitry in V1 encodes distinct forms of binocular conflict and provide a foundation for future studies of both normal binocular function and its disruption in developmental conditions such as strabismus and amblyopia.

## MATERIALS AND METHODS

### Resources availability

#### Lead Contact

Further information and requests for resources and reagents should be directed to and will be fulfilled by the Lead Contact, Eric D. Gaier. Key Resources are outlined in **Supplemental Table 1**.

#### Materials Availability

This study did not generate new unique reagents.

#### Data and Code Availability

The code used to analyze visually evoked potentials are available on Github (https://github.com/danielmontgomery7/suppression-analysis-code). The datasets generated during this study have not been deposited in a public repository but are available from the corresponding author on request.

### Experimental Design and Subject Details

All procedures adhered to the guidelines of the National Institutes of Health and were approved by the Committee on Animal Care at Massachusetts Institute of Technology (MIT). For local field potential and laminar probe experiments, we used male and female mice on a C57BL/6N background (Charles River Laboratories). For principal cell calcium imaging experiments, EMX1.Cre mice (B6.129S2-Emx1tm1(cre)Krj/J, The Jackson Laboratory, RRID:IMSR_JAX:005628) and Ai93(TITL-GCaMP6f)-D;CaMK2a-tTA mice (Igs7tm93.1(tetO−GCaMP6f)Hze Tg(Camk2a-tTA)1Mmay/J, The Jackson Laboratory, RRID:IMSR_JAX:024108) were crossbred in the animal facility at MIT to produce EMX1.GCaMP6f.tTA triple transgenic mice. All EMX1.GCaMP6f.tTA mice were fed with a sterile doxycycline food pellet diet (200 mg/kg, Bio-Serv, Flemington, NJ, USA) until weaned and changed to a normal diet. For SOM+ cell calcium imaging experiments, SOM-Cre mice (B6N.Cg-Ssttm2.1(cre)Zjh/J; catalog #018973, The Jackson Laboratory; RRID:IMSR_JAX:018973). The effects reported in this study did not differ qualitatively by sex, so both were combined. Animals were housed in groups of 2–5 same-sex littermates after weaning at postnatal day 21 (P21). They had access to food and water ad libitum and were maintained on a 12 h light-dark cycle.

### Method Details

#### Surgeries

For local field potential experiments, young adult C57BL/6 mice (P31-32) were injected with 0.1 mg/kg buprenex HCl and subcutaneously (s.c.) to provide analgesia. Induction of anesthesia was achieved via inhalation of isoflurane (3% in oxygen) and thereafter maintained via inhalant isoflurane (1-2% in oxygen). Before surgical incision, the head was shaved and the scalp cleaned with povidone–iodine (10% w/v) and ethanol (70% v/v). The scalp was resected, and the skull surface was scored with a scalpel. A steel head post was affixed to the skull (anterior to bregma) with cyanoacrylate glue. Small burr holes were drilled above both hemispheres of binocular V1 (3.1mm lateral of lambda). Tapered 300–500 kΩ tungsten recording electrodes (FHC), 75µm in diameter at their widest point, were implanted in each hemisphere, 470mm below the cortical surface. Silver wire (A-M Systems) reference electrodes were placed over the left frontal cortex. Electrodes were secured using cyanoacrylate, and the skull was covered with dental cement. Nonsteroidal anti-inflammatory drugs were administered on return to the home cage (meloxicam, 1.0 mg/kg s.c.). Buprenex was administered on post-operative day 1, and meloxicam on post-operative days 1 and 2. Signs of infection and discomfort were carefully monitored. Mice were allowed to recover for at least 48h before head fixation.

For headplate surgeries for acute electrophysiology experiments, adult mice (P60-P80) were anesthetized and prepared as described above. Following scalp incision, a lidocaine (1%) solution was applied onto the periosteum, and the exposed area of skull gently scraped with a scalpel blade. A mark on the skull was made above bV1 (AP −3.6, ML +3.1) with marker. A custom stainless-steel head plate was attached to the skull and dental acrylic (C&B Metabond Quick Adhesive Cement System) was used to form a 5mm x 5mm well above bV1. The well was filled with Kwik-Sil (World Precision Instruments) which was held in place using a thin bridge of Ortho-Jet dental cement (Lang Dental). Mice were returned to their home cage and monitored for at least 48h prior to head fixation. On the morning of the acute recording, mice were placed again under isoflurane anesthesia. The Ortho-Jet bridge and Kwik-Sil cover were removed, and a ∼2mm craniotomy was made around the mark previously made above bV1. Kwik-Sil was once again applied, and the mice were removed from anesthesia and single-house for 2 hours. The mice were head fixed, the Kwik-Sil removed, and a 64-channel laminar probe (H3, Cambridge NeuroTech) was inserted slowly (∼100µm/min) into bV1 perpendicular to the cortical surface. Recordings were obtained and the mice were euthanized immediately thereafter.

For cranial window implantations for two-photon calcium imaging, adult EMX1.GCaMP6f.tTA and SOM-Cre mice (P60-P80) were anesthetized and prepared as described above. A 3mm craniotomy was made over bV1. For SOM-Cre mice, an adeno-associated virus containing the GCaMP7f gene (pGP-AAV9-syn-FLEX-jGCaMP7f-WPRE; catalog #104488-AAV9, Addgene) was loaded into a glass micropipette with a tip diameter of 40–50µm attached to a Nanoject II injection system (Drummond Scientific). The micropipette was then inserted into bV1 (AP −3.6, ML +3.1) layer 2/3 at depths of 300 and 200mm below the pial surface, and 50nL of virus was delivered at each depth. For EMX1.GCaMP6f.tTA mice with genetic expression of GCaMP, an image of the brain was taken during the surgery with a pin at the coordinates for bV1 (AP −3.6, ML +3.1). On subsequent imaging days, the imaging field of view was matched to the vasculature at the coordinates indicated by the pin to ensure that imaging was centered on bV1. Next, a sterile 3-mm-round glass coverslip (CS-3R-0; Warner Instruments) was gently laid on top of the exposed dura mater. The coverslip was secured with cyanoacrylate glue, and a custom stainless steel headplate was attached to the skull. Once the glue had set, dental acrylic (C&B Metabond Quick Adhesive Cement System) was mixed and applied throughout the exposed skull surface. Nonsteroidal anti-inflammatory drugs were administered upon return to the home cage (meloxicam, 1 mg/kg s.c.). Signs of infection and discomfort were carefully monitored. Mice were allowed to recover for at least 3 weeks prior to head-fixation to allow time for the window to clear and for the virus to be fully expressed.

#### Visual stimulus delivery

Prior to stimulus delivery, mice were acclimated to head restraint in front of a gray screen for a 30-minute session on each of two consecutive days. For electrophysiological experiments, mice viewed stimuli on a passive 3D monitor (Vizeo E3D420VX, 120 Hz refresh rate) with reciprocally polarized lenses placed in front of each eye. For calcium imaging experiments, mice viewed stimuli on an active 3D monitor (Acer UMFG6AAB01, 144 Hz refresh rate) with shutter lenses (modified from NVIDIA 3D Vision 2 glasses) placed in front of each eye. Visual stimuli were generated using custom software written in either C++ for interaction with a VSG2/2 card (Cambridge Research Systems) or MATLAB (MathWorks) using the Psychtoolbox extension (http://psychtoolbox.org) to control stimulus drawing and timing and to present different stimuli to each eye. Grating stimuli spanned the full range of monitor display values between black and white, with gamma correction to ensure constant total luminance in both gray-screen and patterned stimulus conditions.

During recordings, mice were head fixed 25 cm from the screen and viewed grating stimuli with a Gabor filter to limit the size of the stimuli to a 40° field of view directly in front of each mouse. Except where noted otherwise, phase-reversing grating stimuli were presented at 1 Hz, 0.05 cycles per degree, and an orientation of 45° or 135°. Stimuli were always presented to the left eye (contralateral to electrode or window) at 100% contrast.

We observed ∼10% interocular bleed-through on both 3D display systems, such that each eye received approximately 10% of the stimulus presented to the other eye. For VEP and 2-photon calcium imaging experiments (**Figures 1-3, 8**), this bleed-through was not corrected. However, the impact on our results was likely minimal given that (1) the ipsilateral eye stimulus was relatively low contrast (10-50%), and (2) ipsilateral eye stimulation generally evokes much weaker responses in mouse V1 compared to the contralateral eye. Thus, any contribution of bleed-through was negligible in the context of our measurements. For acute laminar recordings (**Figures 4-7**), we implemented a correction for this crosstalk by reducing the stimulus presented to each eye by 10% of the contrast intended for the opposite eye. This reduced interocular bleed-through to below detectable levels by eye but resulted in a slightly reduced maximum effective contrast range (0-95% instead of 0-100%).

For VEP recordings, stimuli were presented with the contrast of the right (ipsilateral) eye stimulus increasing from 0% to 50% or decreasing from 50% to 0% in increments of 10% contrast between blocks. Each block lasted 75 seconds with a 15 second interblock interval for 4 blocks per condition. For acute laminar recordings and calcium imaging experiments the right (ipsilateral) eye stimulus was presented at 50% contrast (for Concordant, Phase Offset, and Orthogonal conditions) or 0% contrast (for the Monocular condition). Stimuli for acute laminar recordings were presented in pseudorandomly interleaved 100 second blocks with 30 second interblock intervals for 5 blocks per condition, and stimuli for calcium imaging experiments were presented in pseudorandomly interleaved 10 second blocks with 10 second interblock intervals for 24 blocks per condition.

#### Electrophysiology recordings and analysis

Electrophysiological recordings were conducted in awake, head-restrained mice. Recordings were amplified and digitized using the Recorder-64 system (Plexon Inc.) or the RHD Recording system (Intan Technologies). Local field potentials were recorded from V1 with 1-kHz sampling using a 500-Hz low-pass filter. For the laminar recordings on the Intan system, we sampled at 25 kHz and used a 0.1-Hz high-pass and a 7.5-kHz low-pass filter.

All analyses were conducted using custom MATLAB code and the Chronux toolbox (Bokil, Andrews et al. 2010). For laminar recordings, the raw 25-kHz data from each channel were extracted and converted to mV. Then they were down sampled to 1000Hz and a third order 1- to 300-Hz Butterworth filter was applied. For all data, the mean of the entire channel’s data was subtracted from each time point to account for any DC offset in the system. Next, the data were locally detrended using the locdetrend function in the Chronux toolbox using a 0.5 s window sliding in chunks of 0.1 s. Finally, a third-order Butterworth filter was used to notch frequencies between 58 and 62Hz. For the multiunit activity of laminar recordings, the raw 25-kHz data were extracted for each channel. A 60Hz, 10-dB bandwidth IIR notch filter was applied to each channel and the median value of each channel was subtracted from that channel. The median value across channels for each time point was subtracted from all channel’s timepoints. Visually evoked potentials were normalized by subtracting the average of the first 10 ms of each trial from that trial. A 10-ms moving Gaussian was applied to smooth the visually evoked potential waveform.

#### Current-source density analysis and laminar identification

Current-source density (CSD) analysis on laminar local field potential (LFP) data in V1 was used to identify layer 4 (L4) for each mouse and align our recordings (Mitzdorf 1985, Mitzdorf 1987, Aizenman, Kirkwood et al. 1996). The LFP data were temporally smoothed with a 20 ms Gaussian window, then a five-point hamming window was used to compute the CSD (Ulbert, Halgren et al. 2001). We define 0 mm as the site of the earliest and deepest sink immediately below the superficial source. All other channels were referenced according to that landmark. Thus, superficial channels were above 0 mm and deep channels were below 0 mm. For the sake of reporting in Results, we broke the laminar data into four segments roughly corresponding to each cortical layer (Senzai, Fernandez-Ruiz et al. 2019): L2/3 between +300 and +60 µm, L4 between +60 and −80 µm, L5 between −80 and - 260 µm, and L6 between −260 and −460 µm.

#### Multiunit activity analysis

To measure multiunit activity, we calculated the multiunit activity envelope (MUAe), which provides an instantaneous measure of the number and size of action potentials of neurons in the vicinity of the electrode tip without the requirement of setting an arbitrary threshold level (Legatt, Arezzo et al. 1980, Brosch, Bauer et al. 1995, Super and Roelfsema 2005). MUAe was calculated by applying a third order bandpass (500-5000Hz) Butterworth filter to the common median referenced data to isolate spiking unit activity. Next, the absolute values of the data were taken (units of mV) to remove contamination from far-field signals. Finally, a third order low pass (<250Hz) Butterworth filter was applied to smooth the signal and the data were down sampled to 1000Hz to align with local field potential data. MUAe was z-scored based on the average and standard deviation across all interblock intervals.

#### Single unit activity analysis

Single unit activity was calculated for the common median referenced data using Kilosort 2.5 (Pachitariu, Sridhar et al. 2024) (Kilosort parameters: final projection threshold = 4; spike threshold = 6; channel count = 64; clusters per channel = 4; min spike rate = 0.02; high pass filter frequency = 300; optimization projection threshold = 10; number of blocks = 5; sigma mask = 30; batch size = 1024; whitening range = 32). Spikes identified by Kilosort were manually curated using Phy to isolate single units. Single units with a trough-to-peak latency of less than 0.4 ms were classified as fast-spiking, and units with a latency of greater than 0.4 ms were classified as regular-spiking. The spike rate (in spikes/second) for each unit was calculated by binning spikes into 1 ms time bins for each trial, and then multiplying the number of spikes within each bin across all 500 trials for each condition by 2. Average RS and FS unit PSTHs were calculated by averaging the spike rates of each class of unit. Firing rates during early and late time bins were calculated by averaging firing rates between 40-80 ms after stimulus onset, and 100-200 ms after stimulus onset, respectively. Z-scored SUA was calculated based on the average and standard deviation of the firing rate of each unit across all interblock intervals.

#### In vivo two-photon calcium imaging

Three to four weeks following craniotomy surgery, mice were habituated to the behavior restraint apparatus in front of a gray screen with the objective lens of the two-photon microscope positioned on the head plate for 30 min for two consecutive days before beginning their visual stimulus delivery. A Ti:sapphire laser (Coherent) was used for imaging at a wave length of 930 nm. Photomultiplier tubes (Hamamatsu) and the objective lens (20×, 0.95 numerical aperture, XLUMPLFLN, Olympus) were used to detect fluorescence images. Calcium image recordings were triggered by time-locked transistor-transistor logic pulses generated from the USB-1208fs data acquisition device (Measurement Computing) using PrairieView and TriggerSync software (Bruker) and imaged at a frequency of ∼14.9 Hz at a depth of ∼200 mm in bV1. The size of the imaging field of view was ∼600 × 600 mm^2^ at 512 × 512 pixels.

#### Calcium imaging analysis

Acquired time series of calcium imaging files were processed using Suite2p (Pachitariu, Stringer et al. 2017). All recorded files were registered to stabilize shifts due to animal movement. Regions of interest (ROIs) were automatically detected, then manually curated based on the maximum projection of all frames. Mice were excluded from all further analysis if they had fewer than 100 ROIs within the field of view, as this indicated excessive noise during the recording. We extracted the fluorescence of each ROI for all time points. In line with previous work, for each ROI we calculated the estimated true fluorescence of the ROI. This is the measured fluorescence of the ROI minus 7/10ths of the average measured fluorescence of the surrounding neuropil (Chen, Wardill et al. 2013). We used the average interblock gray period to compute the response relative to gray (F_stim_–F_avg_gray_)/F_avg_gray_. The average population response for each mouse was calculated by taking the average dF/F of all ROIs across all blocks.

### Quantification And Statistical Analysis

#### Statistics

Throughout the results section, all data is expressed as group mean ± standard error of the mean (SEM). Each dataset was assessed for normality and homogeneity of variance prior to choosing a statistical approach, using Levene’s, D’Agostino and Pearson, and, in the case of small sample sizes, Shapiro-Wilk tests. One-way repeated-measures (RM) analysis-of-variance (ANOVA) were used to compare group responses across conditions. Two-way RM ANOVA were used to compare group responses across conditions, stimulus contrasts, or cortical layers. Greenhouse-Geisser was used to modify the degrees of freedom of repeated-measures tests to correct for violations of sphericity. Interactions were followed by tests of simple main effects. Tukey or Sidak’s methods were applied to compensate for multiple comparisons. Uncorrected alpha was set to 0.05. Statistical analyses were performed with Prism 10 (GraphPad; RRID:SCR_002798).

## Acknowledgements

Financial Support: NIH R01 EY023037 (MFB), NIH K08 EY030164 (EDG), Massachusetts Lions Eye Research Fund (EDG)

---

**Figure S1.**
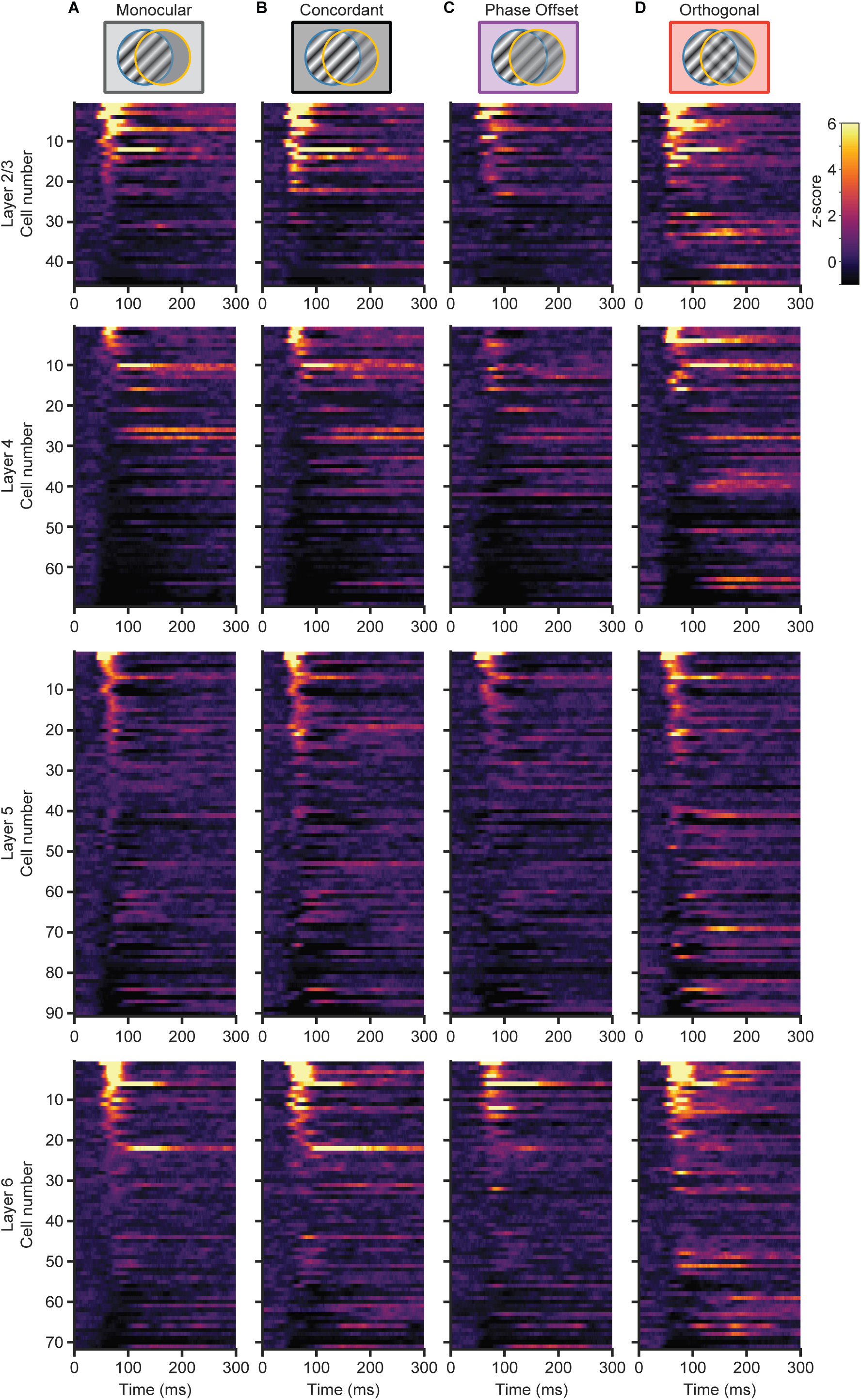
Regular-spiking single unit responses to different stimulus conditions. Z-scored raster plots of regular-spiking (RS) single units for Monocular **(A)**, Concordant **(B)**, Phase Offset **(C)**, and Orthogonal **(D)** stimuli. Units were assigned to L2/3, L4, L5, or L6 based on the location of the electrode contact with the maximal single unit waveform, then sorted based on their average activity between 40-80ms in the Monocular condition. Differences between conditions and layers are quantified in Figure 5.

**Figure S2.**
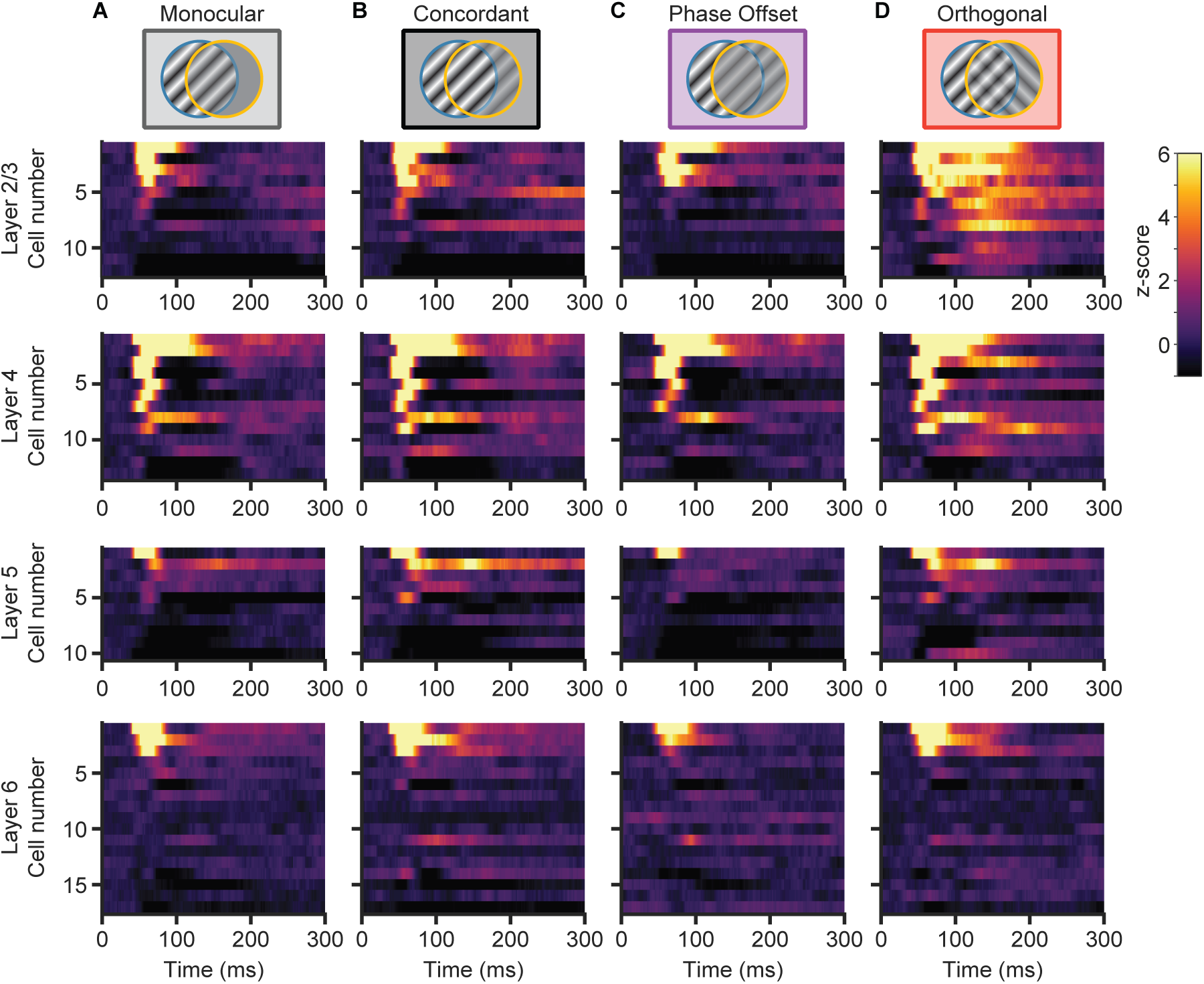
Fast-spiking single unit responses to different stimulus conditions. Same as Figure S3, but for fast-spiking (FS) single units.

**Supplemental Table 1:** Key Resources.

| Reagent or resource | Source | Identifier |
| --- | --- | --- |
| *Experimental models: organisms/strains* |  |  |
| Mouse: C57BL/6N | Bred in lab;<br>Originally from Charles River | RRID:MGI:5651595 |
| Mouse: Emx1-Cre | Bred in lab;<br>Originally from Jackson | RRID:IMSR_JAX:005628 |
| Mouse: Ai93(TITL-GCaMP6f)-D;CaMK2a-tTA | Bred in lab;<br>Originally from Jackson | RRID:IMSR_JAX:024108 |
| Mouse: SOM-IRES-Cre (C57BL/6N) | Bred in lab;<br>Originally from Jackson | RRID:IMSR_JAX:018973 |
| *Software and algorithms* |  |  |
| Raw data | This paper |  |
| MATLAB 2023b | Mathworks | RRID:SCR_001622 |
| Prism 10 | Graphpad | RRID:SCR_002798 |
| Kilosort 2.5 | Github | https://github.com/MouseLand/Kilosort |
| Phy | Github | https://github.com/cortex-lab/phy |
| VEPStimulusSuite:<br>visual stimulus<br>generation and<br>presentation<br>suite | Github (Jeff<br>Gavornik,<br>Boston<br>University) | https://github.com/jeffgavornik/VEPStimulusSuite |
| Analyses were<br>performed using<br>custom-written<br>MATLAB scripts | This paper | https://github.com/danielmontgomery7/suppression-analysis-code |

